# xia2.multiplex: a multi-crystal data analysis pipeline

**DOI:** 10.1101/2022.01.17.476589

**Authors:** Richard J. Gildea, James Beilsten-Edmands, Danny Axford, Sam Horrell, Pierre Aller, James Sandy, Juan Sanchez-Weatherby, C. David Owen, Petra Lukacik, Claire Strain-Damerell, Robin L. Owen, Martin A. Walsh, Graeme Winter

**Affiliations:** Diamond Light Source Ltd, Diamond House, Harwell Science and Innovation Campus, Didcot, Oxfordshire, OX11 0DE, UK; Research Complex at Harwell, Harwell Science and Innovation Campus, Didcot, OX11 0FA, UK

## Abstract

In macromolecular crystallography radiation damage limits the amount of data that can be collected from a single crystal. It is often necessary to merge data sets from multiple crystals, for example small-wedge data collections on micro-crystals, *in situ* room-temperature data collections, and collection from membrane proteins in lipidic mesophase. Whilst indexing and integration of individual data sets may be relatively straightforward with existing software, merging multiple data sets from small wedges presents new challenges. Identification of a consensus symmetry can be problematic, particularly in the presence of a potential indexing ambiguity. Furthermore, the presence of non-isomorphous or poor-quality data sets may reduce the overall quality of the final merged data set.

To facilitate and help optimise the scaling and merging of multiple data sets, we developed a new program, `xia2.multiplex`, which takes data sets individually integrated with *DIALS* and performs symmetry analysis, scaling and merging of multicrystal data sets. `xia2.multiplex` also performs analysis of various pathologies that typically affect multi-crystal data sets, including non-isomorphism, radiation damage and preferential orientation. After describing a number of use cases, we demonstrate the benefit of `xia2.multiplex` within a wider autoprocessing framework in facilitating a multi-crystal experiment collected as part of *in situ* room-temperature fragment screening experiments on the SARS-CoV-2 main protease.

## 1. Introduction

Macromolecular structure determination routinely uses data sets obtained under cryogenic conditions from a single crystal. However, radiation damage limits the amount of data that can be collected from a single crystal. Cryocooling vastly increases the dose that can be tolerated by a single crystal, leading to the dominance of cryocrystallography in macromolecular structure determination (Garman, 1999; Garman & Owen, 2007). However, it is often still necessary to merge multiple data sets from one or more crystals when dealing with radiation sensitive samples and high brilliance X-rays from third generation light sources.

Multi-crystal data collection dates back to the early days of macromolecular crystallography (Kendrew *et al*., 1960; Clemons Jr *et al*., 2001), but has seen a resurgence in recent years (Yamamoto *et al*., 2017) as many scientifically important targets, such as membrane proteins and viruses frequently yield small, weakly diffracting microcrystals. The development of crystallisation in lipidic mesophases (Caffrey, 2003; Caffrey, 2015) and the availability of microfocus beamlines (Evans *et al*., 2011; Smith *et al*., 2012) have facilitated data collection and structure solution of these difficult targets. Data collection strategies for small weakly diffraction crystals rely on collecting many small wedges of data, typically 5-10°per crystal, at cryogenic temperatures. For samples in lipidic mesophase this is often preceded by X-ray raster scanning to identify the location of crystals (Cherezov *et al*., 2007; Rasmussen *et al*., 2011; Rosenbaum *et al*., 2011; Cherezov *et al*., 2009; Warren *et al*., 2013). Such experiments are becoming increasingly automated thanks to developments such as *MeshAndCollect* (Zander *et al*., 2015) and *ZOO* (Hirata *et al*., 2019).

Multi-crystal data collections have also been applied to experimental phasing, where combining data from multiple crystals enhances weak anomalous signals using high-multiplicity data of sufficient quality to enable structure solution by single-wavelength anomalous dispersion (SAD) (Liu *et al*., 2011; Liu & Hendrickson, 2015) and sulfur SAD (S-SAD) (Akey *et al*., 2014; Liu *et al*., 2014; Huang *et al*., 2015; Huang *et al*., 2016; Olieric *et al*., 2016).

Although cryogenic structures have provided the gold standard for structural analysis of macromolecules for decades, it has been shown that cryocooling can hide biologically-significant structural features (Fraser *et al*., 2009; Fraser *et al*., 2011; Fischer *et al*., 2015). Certain classes of macromolecular crystals, such as viruses, can also suffer when cryo-cooled. However, room-temperature data collection presents its own challenges, namely that radiation damage occurs at an absorbed dose one to two orders of magnitude lower than at cryogenic temperatures (Helliwell, 1988; Nave & Garman, 2005). In contrast to cryogenic data collections, an inverse dose-rate effect on crystal lifetime has been observed in room-temperature data (Southworth-Davies *et al*., 2007). As a result, obtaining a complete room-temperature data set from a single crystal is difficult, so combining data from multiple crystals becomes necessary. As demand for room-temperature methods has increased, beamline developments have enabled routine room-temperature data collection on crystals directly from crys-tallisation plates (*in situ*). This has the added benefit of eliminating the need for crystal harvesting (Axford *et al*., 2012; Aller *et al*., 2015; Axford *et al*., 2015), and there now exists a beamline, VMXi at Diamond Light Source, dedicated to *in situ* data collection (Sanchez-Weatherby *et al*., 2019). Advances in beamline and detector technology have enabled the collection of room-temperature data at a higher dose rate (Owen *et al*., 2012; Owen *et al*., 2014; Schubert *et al*., 2016), increasing the general applicability of room-temperature data collection (Aller *et al*., 2015; Broecker *et al*., 2018).

Merging multiple data sets from small wedges presents a number of challenges. For novel structures with unknown space group and unit cell parameters, identifying a consensus symmetry can be problematic, particularly in the presence of indexing ambiguities (Brehm & Diederichs, 2014; Kabsch, 2014; Gildea & Winter, 2018). The presence of non-isomorphous or poor-quality data sets may also degrade the overall quality of the merged data set. Various methods have been developed to identify individual non-isomorphous data sets based on comparison of unit cell parameters (Foadi *et al*., 2013; Zeldin *et al*., 2015) or intensities (Giordano *et al*., 2012; Santoni *et al*., 2017; Diederichs, 2017) to combat this. Rogue data sets, or even individual bad images, can be identified by algorithms such as the 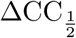 method described by Assmann *et al*. (2016) and implemented within `dials.scale` (Beilsten-Edmands *et al*., 2020).

Microcrystal and room-temperature data collection strategies are a compromise between maximising useful signal, and minimising the effects of radiation damage. By analysing radiation damage we can provide rapid feedback to guide an ongoing experiment and truncate the number of images used to produce the best final composite data set. The *R*_*cp*_ statistic introduced by Winter *et al*. (2019) can also be applied to multi-crystal data, under the assumption that the dose per-image is approximately constant for all data sets. This may be appropriate for multi-crystal data collections where approximately uniformly-sized crystals are bathed in the X-ray beam.

Preferential orientation of crystals can be a concern for some multi-crystal data collections, depending on crystal symmetry and morphology, such as plate-like crystals *in situ* within a flat-bottomed crystallization well. Preferential orientation can lead to under-sampled regions of reciprocal space with systematically low multiplicity or missing reflections, which may have adverse consequences on downstream phasing or refinement. Providing feedback on preferential orientation provides the opportunity for a user to make modifications to their experiment to minimise any resulting issues, for example by fully exploiting the available experimental geometry, or changing the crystallisation conditions or platform (Maeki *et al*., 2016).

Structural biologists have become accustomed to highly automated data analysis provided by synchrotron beamlines around the world (Holton & Alber, 2004; Winter, 2010; Vonrhein *et al*., 2011; Winter & McAuley, 2011; Winter *et al*., 2013; Monaco *et al*., 2013; Yamashita *et al*., 2018), typically obtaining automated data processing results within minutes of the end of data collection for routine experiments. Multi-crystal experiments can generate large volumes of data in minutes, which brings new challenges in terms of bookkeeping and data analysis.

There are two primary aspects in which automated data analysis can support multicrystal experiments. First, rapid feedback from data analysis during beamtime can help guide ongoing experiments, enabling more efficient use of beamtime and allowing a user to more selectively screen sample conditions. Relevant feedback may include suitable metrics on merged data quality, i.e. completeness, multiplicity and resolution, and feedback on experimental pathologies such as non-isomorphism, radiation damage and preferential orientation, that may hinder the experimental goals.

Secondly, after the completion of beamtime, the user may be prepared to invest more time and effort in interactively optimising the best overall data set for any given sample group. Automation is still highly relevant in this context, as the user may have collected data on many sample groups which they wish to process in a similar manner.

Standard autoprocessing pipelines such as *xia2* (Winter, 2010) can handle multicrystal data sets to some extent, however, they are optimised to process a small number of relatively complete data sets, rather than the many tens to hundreds of severely incomplete data sets that comprise a multi-crystal experiment. Recent software developments, for example *KAMO* (Yamashita *et al*., 2018), have focused on automating data processing of multi-crystal experiments.

Here we present new program, `xia2.multiplex`, which has been developed to facilitate the scaling and merging of multiple data sets. It takes as input data sets individually integrated with DIALS and performs symmetry analysis, scaling and merging, and analyses various pathologies that typically affect multi-crystal data sets, including non-isomorphism, radiation damage and preferential orientation. `xia2.multiplex` has been deployed as part of the autoprocessing pipeline at Diamond Light Source, including integration with downstream phasing pipelines such as *DIMPLE* (http://ccp4.github.io/dimple/) and *Big EP* (Sikharulidze *et al*., 2016).

Using data sets collected as part of *in situ* room-temperature fragment screening experiments on the SARS-CoV-2 main protease, we demonstrate the use of `xia2.multiplex` within a wider autoprocessing framework to give rapid feedback during a multi-crystal experiment, and how the program can be used to further improve the quality of final merged data set.

## 2. Methods

Prior to using `xia2.multiplex`, each data set should be processed individually with *DIALS* (Winter *et al*., 2018). Data may be processed either in the primitive, P1, setting, or alternatively Bravais symmetry may be determined prior to integration, using `dials.refine bravais settings`. It is not necessary to individually scale the data at this point.

Preliminary filtering of data sets is performed using hierarchical unit cell clustering methods (Zeldin *et al*., 2015). Histograms and scatterplots of the unit cell distribution are generated for visual analysis, after which symmetry analysis and indexing ambiguity resolution are performed with `dials.cosym`. Finally the data are scaled with `dials.scale`, followed by radiation damage and isomorphism analysis. The main sequence of steps taken by `xia2.multiplex` are outlined in Figure 1.

**Fig. 1.**
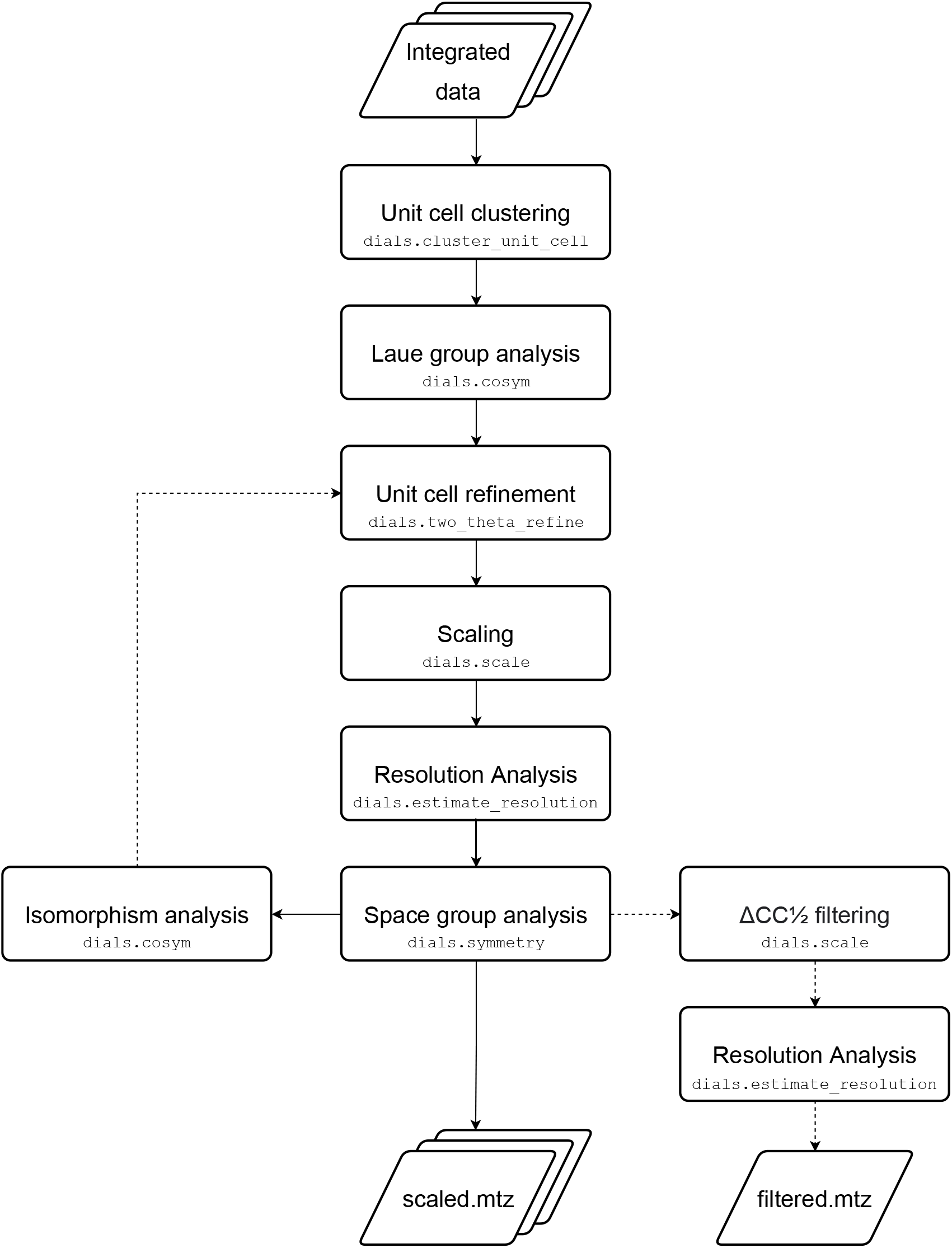
Flowchart outlining the main sequence of steps taken by `xia2.multiplex`. Optional steps are indicated by dashed arrows. The command line programs used at each step are indicated in monospace font.

### 2.1. Symmetry analysis

Initial analysis of the Patterson symmetry of the data is performed using `dials.cosym` (Gildea & Winter, 2018). This is an extension of the methods of Brehm & Diederichs (2014) for resolving indexing ambiguities in partial data sets, for completeness reviewed here.

The maximum possible lattice symmetry compatible with the averaged unit cell is used to compile a list of all potential symmetry operations. The matrix of pairwise correlation coefficients is constructed, of size (*n × m*)^2^, where *n* is the number of data sets and *m* is the number of symmetry operations in the lattice group. The Pearson’s correlation coefficient between data sets *i* and *j*, after application of the *k*th and *l*th symmetry operators respectively, is defined according to

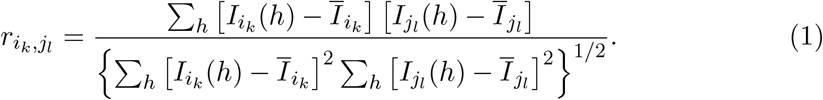

Similarly to Brehm & Diederichs (2014), correlation coefficients are only calculated for pairs of data sets with three or more reflections in common. If a pair of data sets have two or fewer common reflections, then the correlation coefficient for that pair is assumed to be zero. The minimum number of common reflections required for calculation of correlation coefficients is configurable in `dials.cosym` and `xia2.multiplex`. Each data set is represented as *n × m* coordinates in an *m*-dimensional space. Use of an *m*-dimensional space allows the presence of up to *m* orthogonal **x**_*i*_ clusters, where the orthogonality between clusters corresponds to a correlation coefficient 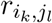 close to zero. A modification of algorithm 2 of Brehm & Diederichs (2014), accounting for the additional symmetry-related copies of each data set, is used to iteratively minimise the function

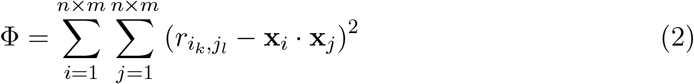

using the L-BFGS minimisation algorithm (Liu & Nocedal, 1989), with randomly-assigned starting coordinates **x** in the range 0–1.

#### 2.1.1. Determination of number of dimensions

It is necessary to use a sufficient number of dimensions to represent any systematic variation present between data sets. Using *m*-dimensional space, where *m* is equal to the number of symmetry operations in the maximum possible lattice symmetry, should be sufficient to represent any systematic variation present due to pseudosymmetry. However, choosing the optimal number of dimensions is a balance between underfitting and overfitting. Using more dimensions than is strictly necessary may reduce the stability of the minimisation, particularly in the case of sparse data, where there is minimal overlap between data sets. As a result, we devised the following procedure to automatically determine the necessary number of dimensions.

1. For each dimension in the range 2–*m* minimise Equation 2 and record the final value of the function.

2. Plot the resulting values as a function of the number of dimensions.

3. Determine the ‘elbow’ point of the plot, in a similar manner to that used by

Zhang *et al*. (2006), to give the optimal number of dimensions.

Alternatively, the user may specify the number of dimensions to be used for the analysis.

#### 2.1.2. Identification of symmetry

A modified form of the algorithms from the program *POINTLESS* (Evans, 2006; Evans, 2011) are used in the determination of the Patterson group symmetry from the results of the initial cosym procedure.

Evans (2011) estimates the likelihood of a symmetry element *S*_*k*_ being present, given the correlation coefficient *CC*_*k*_, as

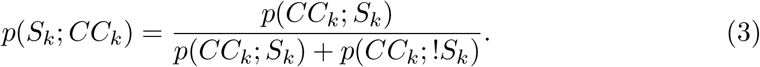

The probability of observing the correlation coefficient *CC*_*k*_ if the symmetry is present, *p*(*CC*_*k*_; *S*_*k*_), is modelled as a truncated Lorentzian centred on the expected value of CC if the symmetry is present, *E*(*CC*; *S*), with a width parameter *γ* = *σ*(*CC*_*k*_).

The distribution of *CC*_*k*_ if the symmetry is not present is modelled as

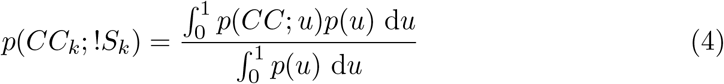

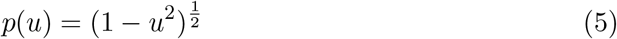

Diederichs (2017) makes clear the relationship between the results of the clustering procedure outlined above, and the correlation coefficient *r*_*ij*_ between two data sets *I* and *j*:

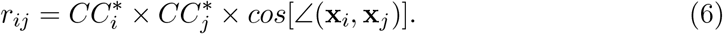

The length of the vectors |**x**_*i*_| are inversely related to the amount of random error, i.e. they provide an estimate of *CC*^*∗*^. The maximum possible correlation coefficient between two data sets is given by the product of their *CC*^*∗*^ values. The angles between two vectors represent genuine systematic differences. For points related by genuine symmetry operations we expect *cos*[∠(**x**_*i*_, **x**_*j*_)] ≈ 1, whereas for points related by symmetry operations that are not present we expect *cos*[∠(**x**_*i*_, **x**_*j*_)] = 0.

We can therefore use *cos*[∠(**x**_*i*_, **x**_*j*_)] in place of *CC*_*k*_, with *E*(*CC*; *S*) = 1. The estimated error *σ*(*CC*_*k*_) used by Evans (2011) has a lower bound of 0.1, which is intended to avoid very small values of *σ* when large numbers of reflections contribute to the calculation of *CC*_*k*_. Since many reflections are contributing indirectly to the angles between any one pair of vectors, we can assume a value of *γ* = 0.1. The average of all observations of *cos*[∠(**x**_*i*_, **x**_*j*_)] corresponding to a given symmetry operator *S*_*k*_, are used as an estimate of *CC*_*k*_.

Once a score has been assigned to each potential symmetry operator, all possible point groups compatible with the lattice group are scored as in Evans (2011) A2:

1. Find the highest lattice symmetry compatible with unit cell dimensions
2. Score each potential rotation operation using all reflections related by that operation
3. Score possible subgroups (Patterson groups) according to combinations of symmetry elements

Once the most likely Patterson group has been identified by the above procedure, it is then relatively straightforward to assign a suitable reindexing operation to each data set to ensure that all data sets are consistently indexed. First, a high density point is chosen as a seed for the cluster. Then, for each data set, identify the nearest symmetry copy of that data set to the seed. The symmetry operation corresponding to this symmetry copy is then the reindexing operation for this data set.

### 2.2. Unit cell refinement

After symmetry determination, an overall best estimate of the unit cell is obtained by refinement of the unit cell parameters against the observed 2*θ* angles, using the program `dials.two theta refine` (Winter *et al*., 2021). This program minimises the unit cell constants against the difference between observed and calculated 2*θ* values, which are determined from background-subtracted integrated centroids. This provides an overall best estimate of the unit cell that is a suitable representative average for use in subsequent downstream phasing and refinement.

### 2.3. Scaling

Data are then scaled using the *physical* scaling model in `dials.scale` (BeilstenEdmands *et al*., 2020). `xia2.multiplex` uses the automatic scaling model selection within `dials.scale` to enable a suitable model parameterisation for both the cases of small-wedge data sets and large-wedge data sets. For small-wedge data sets, each data set is corrected by an overall scale factor and relative B-factor that are smoothly-varying as a function of rotation angle, whereas the absorption correction of the *physical* scaling model is not used as this correction requires the sampling of a diverse set of scattering paths through the sample. For large-wedge data sets, the absorption correction of the *physical* scaling model is used in addition to the smoothly-varying scale and B-factor corrections. The strength of the absorption correction can optionally be set to low (the default), medium or high. This option adjusts the absorption model parameterisation and restraints to enable a correction that more closely matches the expected relative absorption, which can be high at long wavelengths or for crystals containing heavy atoms.

Several rounds of outlier rejection are performed during scaling, to remove individual reflections that have poor agreement with their symmetry-equivalents. The uncertainties of the intensities are also adjusted during scaling, by optimising a single error model across all data sets, in order to account for the effects of systematic errors which tend to increase the variability of intensities within each symmetry-equivalent group. Optionally, for anomalous data, Friedel pairs can be treated separately in scaling, which can increase the strength of the anomalous signal.

#### 2.3.1. Estimation of resolution cutoff

After the data have been successfully scaled, the program `dials.estimate resolution` is used to estimate a suitable resolution cutoff for the data. By default, this is determined from a fit of a hyperbolic tangent to 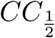 calculated in resolution bins, similar to that used by *AIMLESS* (Evans & Murshudov, 2013). The resolution cutoff is chosen as the resolution where the fit curve reaches 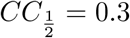 (this cutoff value can be controlled by the user). A second round of scaling with `dials.scale` is then performed after application of the resolution cutoff. The default cutoff value of 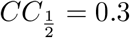 is chosen as one that works well in the context of autoprocessing in order to provide a consistent set of merging statistics for judging data quality during and ongoing experiment. Suitable cutoff values may depend on the downstream data processing requirements, but the current gold standard for publication is to use “paired refinement” to determine the resolution at which including higher resolution data in refinement no longer improves the model (Karplus & Diederichs, 2012).

#### 2.3.2. Space group identification

After the data have been scaled in the Patterson group identified by `dials.cosym` (§2.1.2), analysis of potential systematic absences is performed by `dials.symmetry` in order to assign a final space group. In this analysis, the existence of each potential screw axis allowed by the Patterson group is tested, by calculating the z-score based on the deviation from zero of the merged < *I/σ*(*I*) *>* for the expected absent reflections. From the individual z-scores, a likelihood for the presence of each screw axis is determined, which are combined to score and select the most likely non-enantiogenic space group.

### 2.4. Radiation damage analysis

`xia2.multiplex` performs a number of analyses that can be useful in assessing the extent of any radiation damage which may be present. Plots of scale factor and *R*_*merge*_ vs. image number are generated to look for any trends associated with radiation damage. The *R*_*cp*_ statistic introduced by Winter *et al*. (2019) can also be applied to multi-crystal data. This statistic accumulates the pairwise measured intensity differences as a function of dose (or image number). In the absence of accurate dose information for each data set it is necessary to make the assumption the dose perimage is approximately constant for all data sets. In order to assess how many images per crystal are necessary to achieve a complete data set, a plot of completeness vs. dose is also generated.

### 2.5. Isomorphism analysis

Unit cell clustering, as implemented in the program BLEND (Foadi *et al*., 2013) and elsewhere (Zeldin *et al*., 2015), is used by `xia2.multiplex` as a preliminary filtering step to reject any highly non-isomorphous data sets. `xia2.multiplex` implements two alternative intensity-based clustering methods that are suitable for identification and analysis of non-isomorphism, once symmetry-determination, resolution of indexing ambiguities, and scaling have been carried out as described above. Clustering on correlation coefficients (Giordano *et al*., 2012; Santoni *et al*., 2017; Yamashita *et al*., 2018) begins by calculating a matrix of pairwise correlation coefficients:

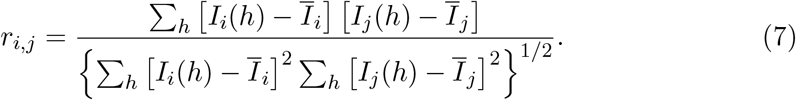

A distance matrix defined as *d*_*i,j*_ = 1 *−r*_*i,j*_ is provided as input to the SciPy (Virtanen *et al*., 2020) hierarchical clustering routine using the *average* linkage method. Clusters are sorted by distance, and the completeness and multiplicity of each cluster is reported. Optionally, `xia2.multiplex` can scale and merge the data sets defined by each cluster that meets user-defined criteria for minimum completeness or multiplicity. A second intensity-based clustering method follows that described by Diederichs (2017) who demonstrated that the methods of Brehm & Diederichs (2014) could be generalised to search for any systematic differences between data sets, not just those caused by an indexing ambiguity. In addition to its use for identifying the Patterson symmetry (§2.1.2), `dials.cosym` can also be used for analysis of non-isomorphism. In this mode, rather than searching for the presence of potential additional symmetry operators, the matrix of pairwise correlation coefficients of size *n*^2^ reduces to Equation 7. The function defined by Equation 2 is minimised as before to obtain a representation of the similarity between data sets in a reduced dimensional space.

As made clear by Diederichs (2017), the length of a vector, **x**_*i*_ is inversely proportional to the random error in data set **X**_*i*_. The angle between vectors **x**_*i*_ and **x**_*j*_ corresponds to the level of systematic error between data sets **X**_*i*_ and **X**_*j*_, and can thus be used to estimate the degree of non-isomorphism between those data sets. Analysis of the angular separation of vectors, **x**, can be used to identify groups of systematically different data sets. Hierarchical clustering on the cosines of the angles between vectors is performed to identify possible groupings of data sets for further investigation. Optionally `xia2.multiplex` can re-scale multiple subsets of data, which can be controlled by specifying a maximum number of clusters to merge and/or the minimum required completeness or multiplicity for a cluster.

The final approach to isomorphism analysis implemented within `xia2.multiplex` is the 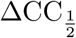 method described by Assmann *et al*. (2016), and implemented within`dials.scale` (Beilsten-Edmands *et al*., 2020). If 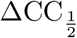 filtering is selected, then `xia2.multiplex` will perform additional scaling with `dials.scale`, reject any data sets that are identified as significant outliers according to 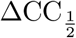 analysis. Whilst this approach may not be suitable if there are two or more significant non-isomorphous populations, it may give useful results if there are a small number of data sets that are systematically different from the majority.

### 2.6. Preferential orientation

The report generated by `xia2.multiplex` includes stereographic projections of crystal orientation relative to the laboratory frame, generated with the program `dials.stereographic projection`. A random distribution of points (each point corresponds to a crystal, or its symmetry equivalent) in a stereographic projection suggests a random distribution of crystal orientation, whereas a systematic non-random distribution may be indicative of preferential crystal orientation. `xia2.multiplex` also generates a number of plots that can aid in the analysis of the distribution of multiplicities.

A new command, `dials.missing reflections`, has been developed to identify connected regions of missing reflections in reciprocal space. This is achieved by first generating the complete set of possible miller indices, then performing connected components analysis on a graph of nearest neighbours in the list of missing reflections, taking into account any symmetry operations that may be present. Prior to performing the analysis, it is necessary to map centred unit cells to the primitive setting, in order to avoid systematically absent reflections complicating the analysis. The complete set of possible miller indices are generated, and expanded to cover the full sphere of reciprocal space by application of symmetry operators belonging to the known space group. This allows the identification of connected regions that cross the boundary of the asymmetric unit. Nearest neighbour analysis is used to construct a graph of connected regions which is then used to perform connected components analysis to identify each connected region of missing reflections. Miller indices for missing reflections are then mapped back to the asymmetric unit in order to identify the set of unique miller indices belonging to each region. A sorted list of connected regions is reported to the user, detailing the resolution range spanned by each region, and the number and proportion of total reflections comprising each each region.

## 3. Deployment of xia2.multiplex at Diamond Light Source

`xia2.multiplex` as described above has been deployed as part of the autoprocessing pipeline at Diamond Light Source. A series of partial data sets are collected from a set of related crystals, for example from multiple crystals within one or more drops in a crystallisation plate (Sanchez-Weatherby *et al*., 2019), sample loop, or sample mesh. After the end of each data collection, the partial data set is processed individually with *DIALS* via *xia2*. On the successful completion of *xia2*, a `xia2.multiplex` processing job is triggered using as input all successful *xia2* results from this and prior data collections. `xia2.multiplex` results, including merging statistics, are recorded in ISPyB (Delagenière *et al*., 2011) for presentation to the user via SynchWeb (Fisher *et al*., 2015), where results are typically available within minutes of the end of the data collection. Prior to data collection, users may define groups of related samples for combining with `xia2.multiplex`, either via SynchWeb or via a configuration file in a pre-defined location. In the absence of this information, `xia2.multiplex` will only combine data collected on the same *sample*, i.e. loop, mesh or well within a crystallisation plate.

If a PDB file has been associated with the data collection, then automated structure refinement is performed with the program *DIMPLE* (http://ccp4.github.io/dimple/) using the merged reflections output by `xia2.multiplex`.

## 4. Examples

### 4.1. Room-temperature in situ experimental phasing

Using data from (Lawrence *et al*., 2020), we showcase the use of `xia2.multiplex` applied to multi-crystal room-temperature *in situ* data sets from heavy-atom soaks of Lysozyme crystals, demonstrating successful experimental phasing using the resulting `xia2.multiplex` output. Data from Lysozyme crystals soaked with six different heavy atom solutions were processed individually with *DIALS* via *xia2* followed by symmetry determination, scaling and merging with `xia2.multiplex`. Partial data sets identified as outliers according to 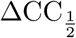 were rejected in an automated iterative process with `xia2.multiplex`. Data processing statistics for each heavy atom soak, with and without 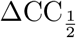 filtering of outlier data sets, are shown in Tables 1 and 2. Phasing was performed with *fast ep* using *SHELXC/D/E* (Sheldrick, 2010). Structure refinement was performed by REFMAC (Murshudov *et al*., 2011) via *DIMPLE* using the reference structure 6QQF (Gotthard *et al*., 2019). Anomalous difference maps were calculated by *ANODE* (Thorn & Sheldrick, 2011) via the `--anode` option in *DIMPLE*. Significant anomalous signal was observed, as indicated by the *SHELXC* plot of < *d*”*/sigI >* vs. resolution (Figure 2a). Substructure searches with *SHELXD* were successful (Figure 2b), and traceable electron-density maps were obtained by *SHELXE*. Anomalous difference maps calculated by *ANODE* (Thorn & Sheldrick, 2011) via *DIMPLE* indicated the presence of significant anomalous difference peaks (Figures 2c and d).

**Table 1.**
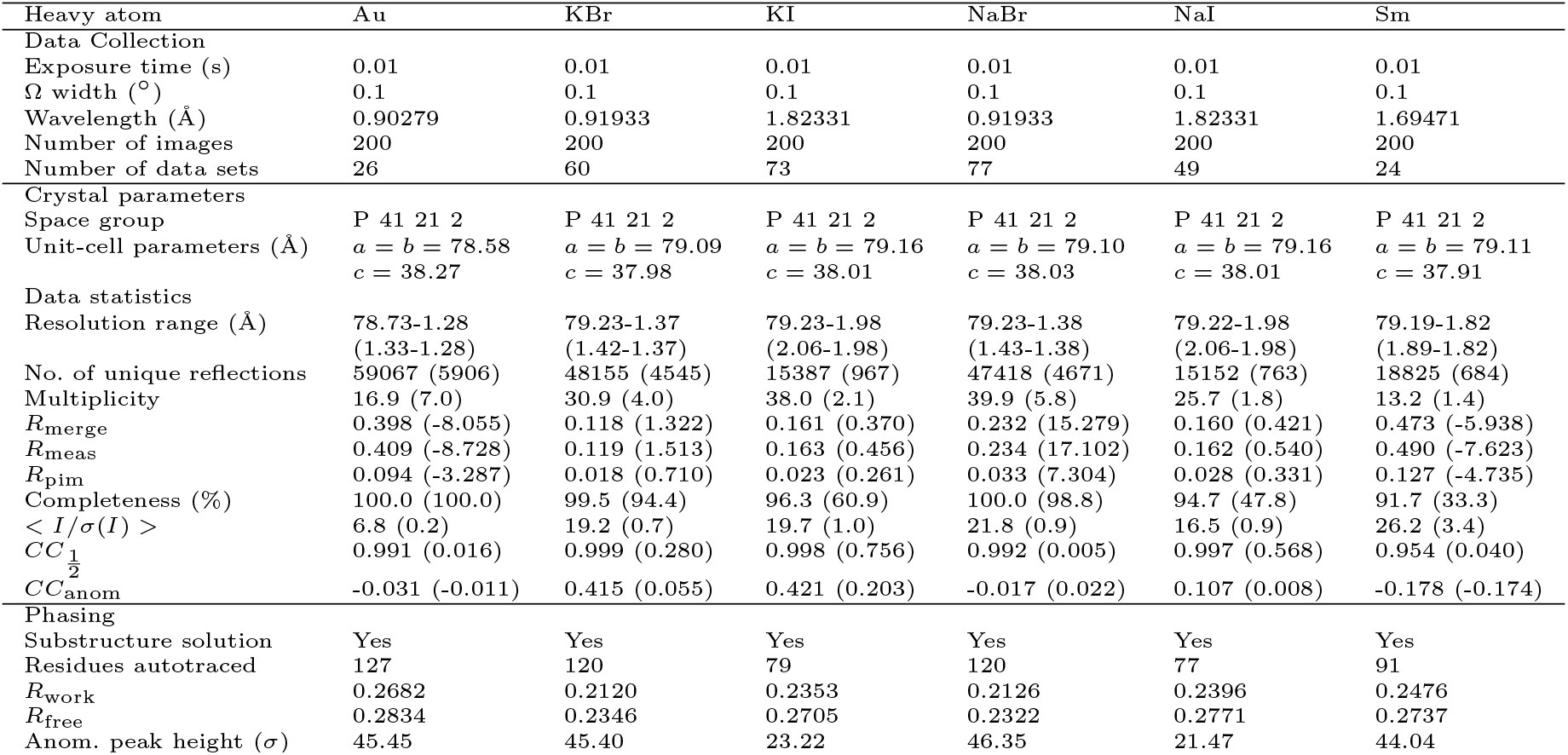
Data collection, merging and refinement statistics for Lysozyme room-temperature in situ heavy atom soaks using all data sets. Values in parentheses are for the highest resolution shell.

| Heavy atom | Au | KBr | KI | NaBr | NaI | Sm |
| --- | --- | --- | --- | --- | --- | --- |
| Data Collection |  |  |  |  |  |  |
| Exposure time (s) | 0.01 | 0.01 | 0.01 | 0.01 | 0.01 | 0.01 |
| $\Omega$ width ($^\circ$) | 0.1 | 0.1 | 0.1 | 0.1 | 0.1 | 0.1 |
| Wavelength ($\text{\AA}$) | 0.90279 | 0.91933 | 1.82331 | 0.91933 | 1.82331 | 1.69471 |
| Number of images | 200 | 200 | 200 | 200 | 200 | 200 |
| Number of data sets | 26 | 60 | 73 | 77 | 49 | 24 |
| Crystal parameters |  |  |  |  |  |  |
| Space group | P 41 21 2 | P 41 21 2 | P 41 21 2 | P 41 21 2 | P 41 21 2 | P 41 21 2 |
| Unit-cell parameters ($\text{\AA}$) | $a = b = 78.58$<br>$c = 38.27$ | $a = b = 79.09$<br>$c = 37.98$ | $a = b = 79.16$<br>$c = 38.01$ | $a = b = 79.10$<br>$c = 38.03$ | $a = b = 79.16$<br>$c = 38.01$ | $a = b = 79.11$<br>$c = 37.91$ |
| Data statistics |  |  |  |  |  |  |
| Resolution range ($\text{\AA}$) | 78.73-1.28<br>(1.33-1.28) | 79.23-1.37<br>(1.42-1.37) | 79.23-1.98<br>(2.06-1.98) | 79.23-1.38<br>(1.43-1.38) | 79.22-1.98<br>(2.06-1.98) | 79.19-1.82<br>(1.89-1.82) |
| No. of unique reflections | 59067 (5906) | 48155 (4545) | 15387 (967) | 47418 (4671) | 15152 (763) | 18825 (684) |
| Multiplicity | 16.9 (7.0) | 30.9 (4.0) | 38.0 (2.1) | 39.9 (5.8) | 25.7 (1.8) | 13.2 (1.4) |
| $R_{\text{merge}}$ | 0.398 (-8.055) | 0.118 (1.322) | 0.161 (0.370) | 0.232 (15.279) | 0.160 (0.421) | 0.473 (-5.938) |
| $R_{\text{meas}}$ | 0.409 (-8.728) | 0.119 (1.513) | 0.163 (0.456) | 0.234 (17.102) | 0.162 (0.540) | 0.490 (-7.623) |
| $R_{\text{pim}}$ | 0.094 (-3.287) | 0.018 (0.710) | 0.023 (0.261) | 0.033 (7.304) | 0.028 (0.331) | 0.127 (-4.735) |
| Completeness (%) | 100.0 (100.0) | 99.5 (94.4) | 96.3 (60.9) | 100.0 (98.8) | 94.7 (47.8) | 91.7 (33.3) |
| $\langle I/\sigma(I) \rangle$ | 6.8 (0.2) | 19.2 (0.7) | 19.7 (1.0) | 21.8 (0.9) | 16.5 (0.9) | 26.2 (3.4) |
| $CC_{\frac{1}{2}}$ | 0.991 (0.016) | 0.999 (0.280) | 0.998 (0.756) | 0.992 (0.005) | 0.997 (0.568) | 0.954 (0.040) |
| $CC_{\text{anom}}$ | -0.031 (-0.011) | 0.415 (0.055) | 0.421 (0.203) | -0.017 (0.022) | 0.107 (0.008) | -0.178 (-0.174) |
| Phasing |  |  |  |  |  |  |
| Substructure solution | Yes | Yes | Yes | Yes | Yes | Yes |
| Residues autotraced | 127 | 120 | 79 | 120 | 77 | 91 |
| $R_{\text{work}}$ | 0.2682 | 0.2120 | 0.2353 | 0.2126 | 0.2396 | 0.2476 |
| $R_{\text{free}}$ | 0.2834 | 0.2346 | 0.2705 | 0.2322 | 0.2771 | 0.2737 |
| Anom. peak height ($\sigma$) | 45.45 | 45.40 | 23.22 | 46.35 | 21.47 | 44.04 |

**Table 2.** Data collection, merging and refinement statistics for Lysozyme room-temperature in situ heavy atom soaks after removal of data sets identified by 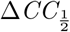 analysis. Values in parentheses are for the highest resolution shell.

| Heavy atom | Au | KBr | KI | NaBr | NaI | Sm |
| --- | --- | --- | --- | --- | --- | --- |
| Data Collection |  |  |  |  |  |  |
| Exposure time (s) | 0.01 | 0.01 | 0.01 | 0.01 | 0.01 | 0.01 |
| $\Omega$ width ($^\circ$) | 0.1 | 0.1 | 0.1 | 0.1 | 0.1 | 0.1 |
| Wavelength ($\text{\AA}$) | 0.90279 | 0.91933 | 1.82331 | 0.91933 | 1.82331 | 1.69471 |
| Number of images | 200 | 200 | 200 | 200 | 200 | 200 |
| Number of data sets | 22 | 59 | 72 | 75 | 48 | 22 |
| Crystal parameters |  |  |  |  |  |  |
| Space group | P 41 21 2 | P 41 21 2 | P 41 21 2 | P 41 21 2 | P 41 21 2 | P 41 21 2 |
| Unit-cell parameters ($\text{\AA}$) | $a = b = 78.58$<br>$c = 38.27$ | $a = b = 79.09$<br>$c = 37.98$ | $a = b = 79.17$<br>$c = 38.01$ | $a = b = 79.10$<br>$c = 38.03$ | $a = b = 79.16$<br>$c = 38.01$ | $a = b = 79.11$<br>$c = 37.91$ |
| Data statistics |  |  |  |  |  |  |
| Resolution range ($\text{\AA}$) | 39.31-1.27<br>(1.32-1.27) | 79.23-1.35<br>(1.40-1.35) | 79.23-1.98<br>(2.06-1.98) | 79.24-1.33<br>(1.38-1.33) | 79.22-1.98<br>(2.06-1.98) | 79.19-1.82<br>(1.89-1.82) |
| No. of unique reflections | 60440 (6006) | 49744 (4213) | 15363 (945) | 51260 (3621) | 15096 (726) | 18509 (589) |
| Multiplicity | 13.1 (5.1) | 29.1 (2.7) | 36.1 (2.0) | 34.4 (2.2) | 24.2 (1.7) | 10.7 (1.3) |
| $R_{\text{merge}}$ | 0.137 (2.345) | 0.115 (1.161) | 0.163 (0.336) | 0.111 (1.106) | 0.156 (0.346) | 0.074 (0.162) |
| $R_{\text{meas}}$ | 0.142 (2.620) | 0.116 (1.390) | 0.165 (0.417) | 0.112 (1.361) | 0.159 (0.440) | 0.077 (0.216) |
| $R_{\text{pim}}$ | 0.036 (1.131) | 0.018 (0.741) | 0.023 (0.241) | 0.015 (0.772) | 0.028 (0.266) | 0.020 (0.141) |
| Completeness (%) | 100.0 (99.6) | 98.4 (83.7) | 96.1 (59.5) | 96.8 (68.5) | 94.4 (45.5) | 90.2 (28.6) |
| $\langle I/\sigma(I) \rangle$ | 7.2 (0.2) | 20.3 (0.8) | 20.1 (1.3) | 21.9 (0.6) | 17.3 (1.4) | 25.3 (3.7) |
| $CC_{\frac{1}{2}}$ | 0.997 (0.187) | 0.999 (0.313) | 0.997 (0.802) | 0.999 (0.315) | 0.994 (0.736) | 0.996 (0.894) |
| $CC_{\text{anom}}$ | 0.313 (0.011) | 0.455 (-0.020) | 0.565 (-0.100) | 0.423 (0.089) | 0.485 (0.055) | 0.656 (0.024) |
| Phasing |  |  |  |  |  |  |
| Substructure solution | Yes | Yes | Yes | Yes | Yes | Yes |
| Residues autotraced | 116 | 103 | 85 | 114 | 54 | 119 |
| $R_{\text{work}}$ | 0.2668 | 0.2116 | 0.2355 | 0.2140 | 0.2420 | 0.2078 |
| $R_{\text{free}}$ | 0.2820 | 0.2333 | 0.2704 | 0.2332 | 0.2753 | 0.2424 |
| Anom. peak height ($\sigma$) | 46.47 | 45.43 | 23.22 | 47.63 | 21.85 | 47.00 |

**Fig. 2.**
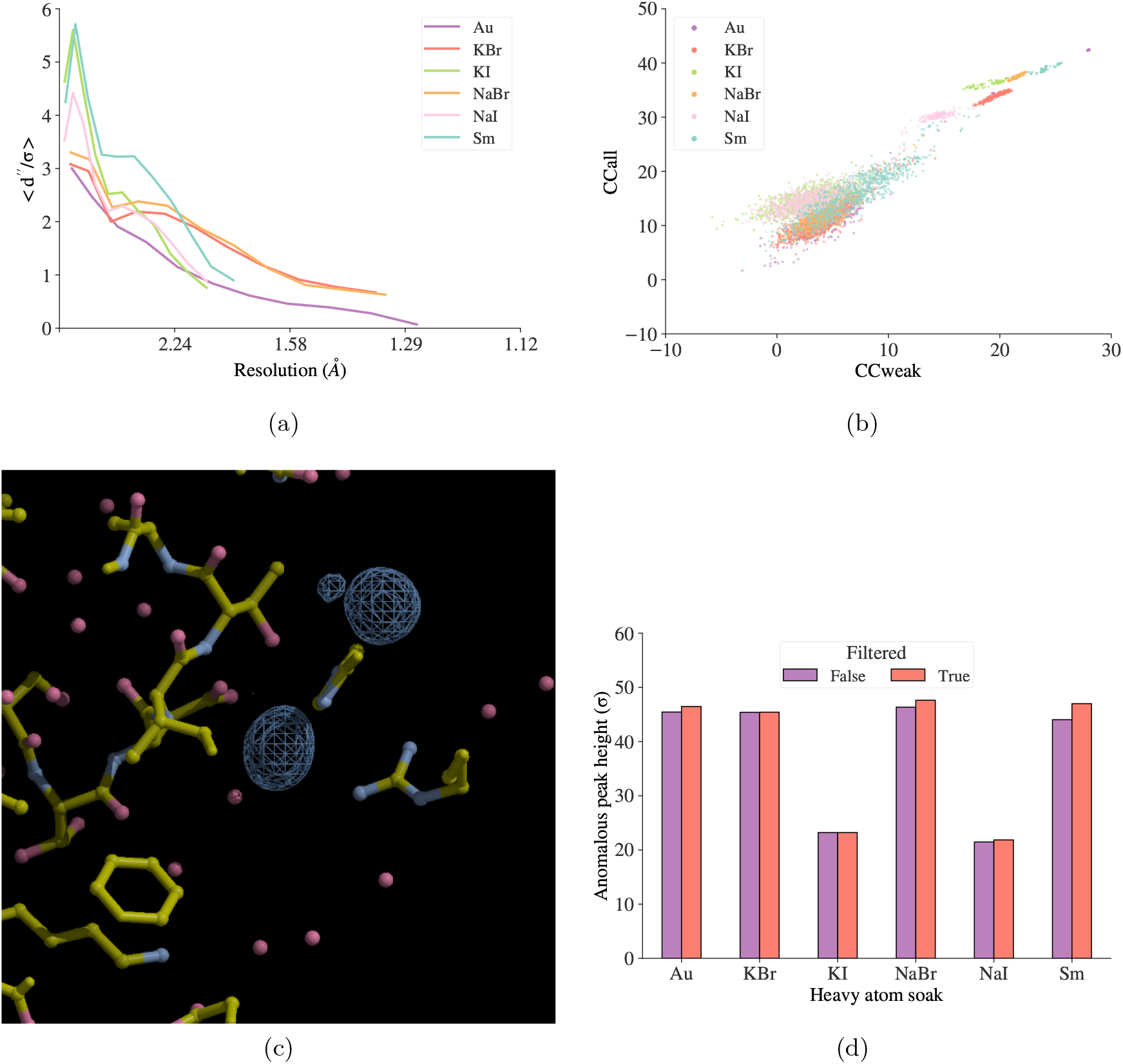
Experimental phasing and anomalous signal from multi-crystal roomtemperature *in situ* experiments using lysozyme crystals soaked with various heavy atom solutions. (a) *SHELXC* plot of < *d*”*/sigI >*. (b) *CC*_all_ vs. *CC*_weak_ after substructure solution with HKL2MAP/SHELXD. (c) Anomalous difference map peaks identified by *ANODE* via *DIMPLE* for lysozyme Au soaks. Contours are drawn at 4*σ*. (d) Anomalous difference map peak heights identified by *ANODE* via *DIMPLE*, with and without filtering of outlier regions of data sets.

To assess the impact of 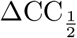 filtering on the resulting anomalous signal, we performed experimental phasing, structure refinement (via *DIMPLE*) and calculated anomalous difference maps using data both with and without 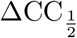 filtering of out-liers. Substructure solution and autotracing were successful in both cases. 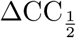 filtering also resulted in improved merging statistics, typically in 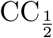, CC_anom_, < *d*”*/sigI* >, < *I/σ*(*I*) > and R_pim_ vs. resolution (Tables 1 and 2). For the NaBr and Sm soaks there is a particularly significant improvement in *R*_work_ and *R*_free_ after 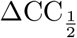 filtering. These two soaks also correspond to the data sets that showed the largest improvement in anomalous difference peak height after removal of outlier data sets (Figure 2d).

We note that merging statistics such as correlation coefficients and R-factors, which are calculated only on the unmerged intensity values without taking into account their errors, can be affected by regions of lower data quality that are suitably downweighted with larger errors during scaling. The presence of these regions however does not adversely affect the resulting merged intensities, which are appropriately weighted. This disparity is most likely to be evident for high multiplicity data with regions of significant radiation damage, in which case merged data quality indicators are most representative of the data quality.

As outlined in §2.5, there are several different methods available in `xia2.multiplex` for identifying outlier data sets. Above, we used 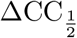 filtering to identify and exclude outlier partial data sets. Visualisation of the distribution and hierarchical clustering on unit cell parameters for the Sm soak (Figure 3e and f) identify data set 11 as an outlier, which was also the first data set to be excluded by 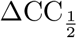 filtering. Similarly, hierarchical clustering on pairwise correlation coefficients (Figure 4a) and on the cosines of the angles between vectors, **x**, (Figure 4b) both identify data set 11 as an outlier. Whilst in this case, all available methods for isomorphism analysis identified data set 11 as the least compatible data set, it is beneficial to have an array of different methods available, as the best method for a particular system may depend on the nature of any isomorphism involved.

**Fig. 3.**
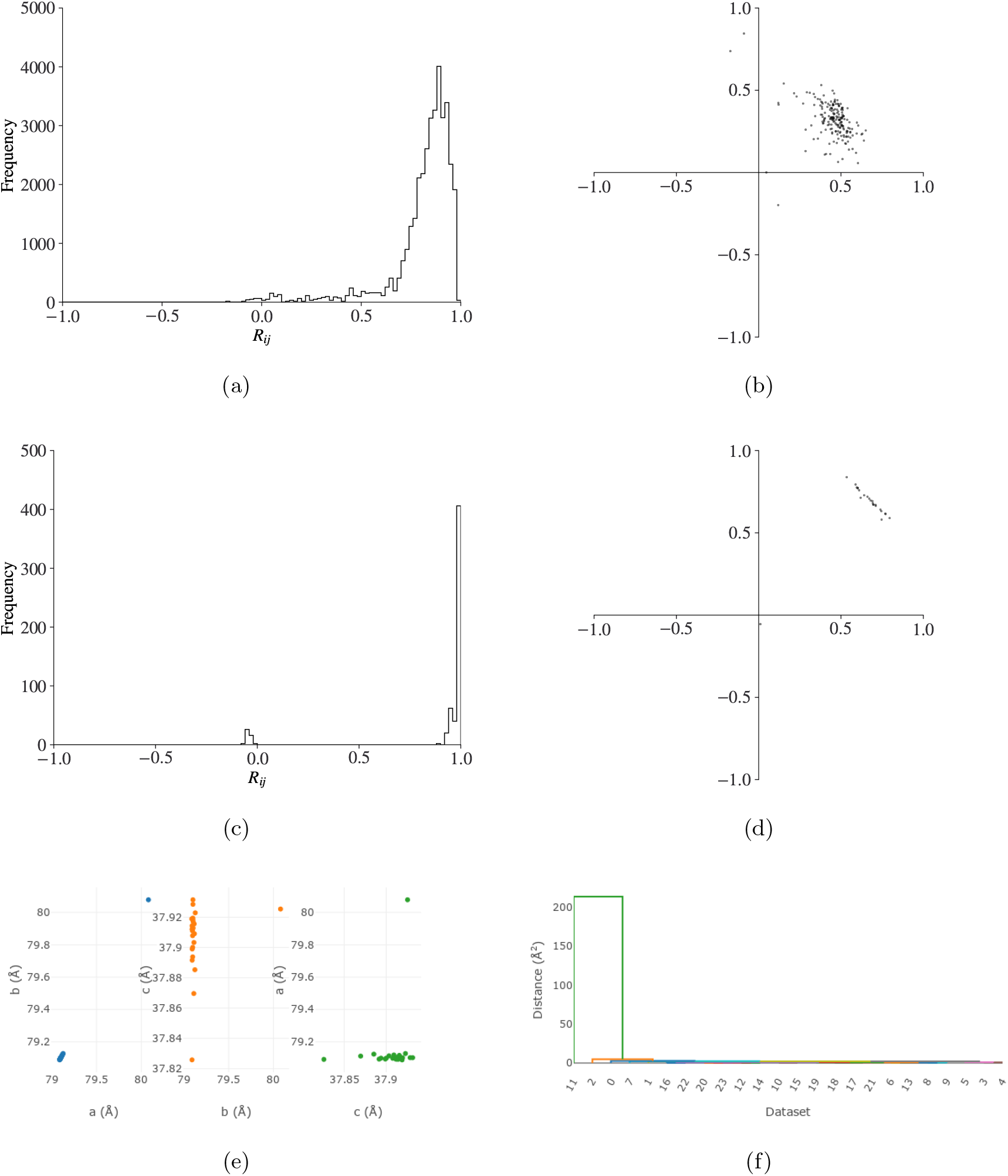
dials.cosym plots for data from Lysozyme Sm soaks as described in §4.1. (a) histogram of (*n × m*)^2^ pairwise *R*_*ij*_ correlation coefficients and (b) the (*n × m*) vectors **x** determined by the minimisation of Equation 2 during symmetry determination with dials.cosym. The *R*_*ij*_ correlation coefficients are clustered towards 1 and the majority of the vectors **x** form a single cluster, suggesting the absence of an indexing ambiguity, i.e. the Patterson group of the data set corresponds to the maximum lattice symmetry. (c) and as above, but after symmetry determination and scaling. The distribution of the *n*^2^ *R*_*ij*_ correlation coefficients is sharpened towards 1 as scaling improves the internal consistency of the data. There is also an effect from multiplicity when comparing to (a), as here the *n*^2^ *R*_*ij*_ values are calculated in the highest symmetry group for the lattice. All but one of the *n* vectors **x** form a tight cluster, with the vector lengths close to 1. Visualisation of the distribution of unit cell parameters (e) and clustering on unit cell parameters (f)suggests the presence of an outlier data set.

**Fig. 4.**
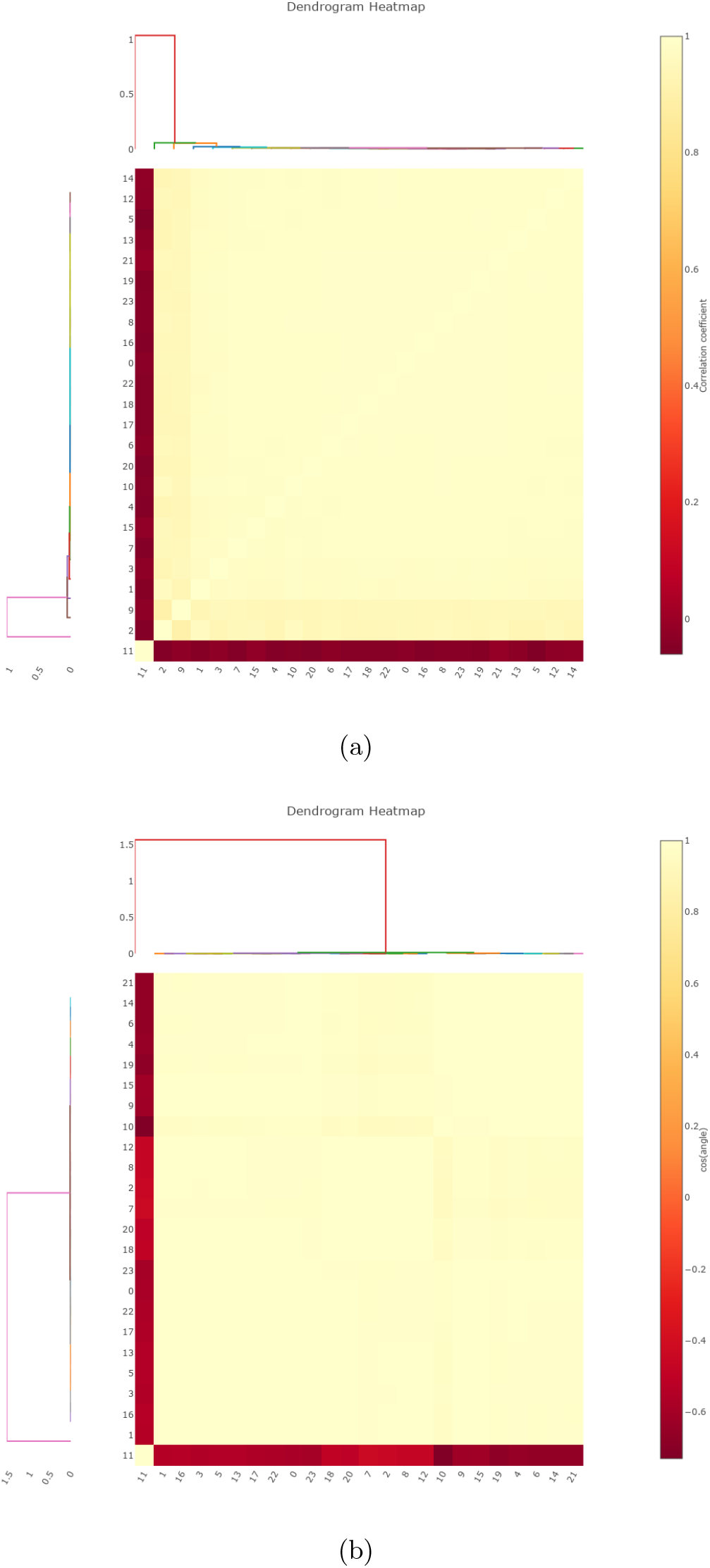
Hierarchical clustering (a) on pairwise correlation coefficients and (b) on the cosines of the angles between vectors in Figure 3d identify the presence of an outlier data set.

### 4.2. TehA

Previously published *in situ* data for Haemophilus influenzae TehA (Axford *et al*., 2015) were used to further demonstrate the applicability of `xia2.multiplex` and the tools contained therein. 73 partial data sets were processed individually with *DIALS* via *xia2*, providing no prior space group or unit cell information. 71 successfully-integrated data sets were provided as input to `xia2.multiplex`, where data were combined and scaled using `dials.cosym` and `dials.scale`. Two data sets were identified as having inconsistent unit cells by preliminary filtering and removed, leaving 69 data sets for subsequent symmetry analysis and scaling. Structure refinement was performed by REFMAC (Murshudov *et al*., 2011) via *DIMPLE*. Data processing and refinement statistics using all data, and only those remaining after filtering by 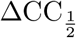, are shown in Table 3.

**Table 3.** Data collection, merging and refinement statistics for TehA. Values in parentheses are for the highest resolution shell

| | All data | $\Delta CC_{\frac{1}{2}}$-filtered data |
| --- | --- | --- |
| Data Collection |  |  |
| Exposure time (s) | 0.04 | 0.04 |
| $\Omega$ width ($^{\circ}$) | 0.2 | 0.2 |
| Transmission (%) | 12.34 | 12.34 |
| Number of images | 20-50 | 20-50 |
| Number of data sets | 69 | 64 |
| Crystal parameters |  |  |
| Space group | R 3 :H | R 3 :H |
| Unit-cell parameters ($\text{\AA}$) | $a = b = 98.76, c = 136.77$ | $a = b = 98.76, c = 136.77$ |
| Data statistics |  |  |
| Resolution range ($\text{\AA}$) | 72.56-2.13 (2.21-2.13) | 72.56-2.14 (2.22-2.14) |
| No. of unique reflections | 26203 (2415) | 25851 (2396) |
| Multiplicity | 13.7 (6.8) | 13.0 (6.7) |
| $R_{\text{merge}}$ | 0.315 (-1033.253) | 0.162 (2.508) |
| $R_{\text{meas}}$ | 0.326 (-1113.925) | 0.167 (2.703) |
| $R_{\text{pim}}$ | 0.078 (-406.346) | 0.040 (0.981) |
| Completeness (%) | 94.1 (86.3) | 94.2 (86.8) |
| $\langle I/\sigma(I) \rangle$ | 13.1 (1.3) | 13.9 (1.5) |
| $CC_{\frac{1}{2}}$ | 0.988 (0.285) | 0.996 (0.360) |
| $CC_{\text{anom}}$ | -0.002 (0.004) | 0.073 (0.045) |
| $R_{\text{work}}$ | 0.1515 | 0.1499 |
| $R_{\text{free}}$ | 0.1726 | 0.1711 |

The maximum possible lattice symmetry was determined to be *R −* 3*m*:*H*, with a maximum of six symmetry operations. Analysis of the value of Equation 2 as a function of the number of dimensions identified that two dimensions were sufficient to explain the variation between data sets. Further symmetry analysis with `dials.cosym` correctly identified the Patterson group as *R −* 3:*H*, resolving the indexing ambiguity present in this space group (Figure 5b).

**Fig. 5.**
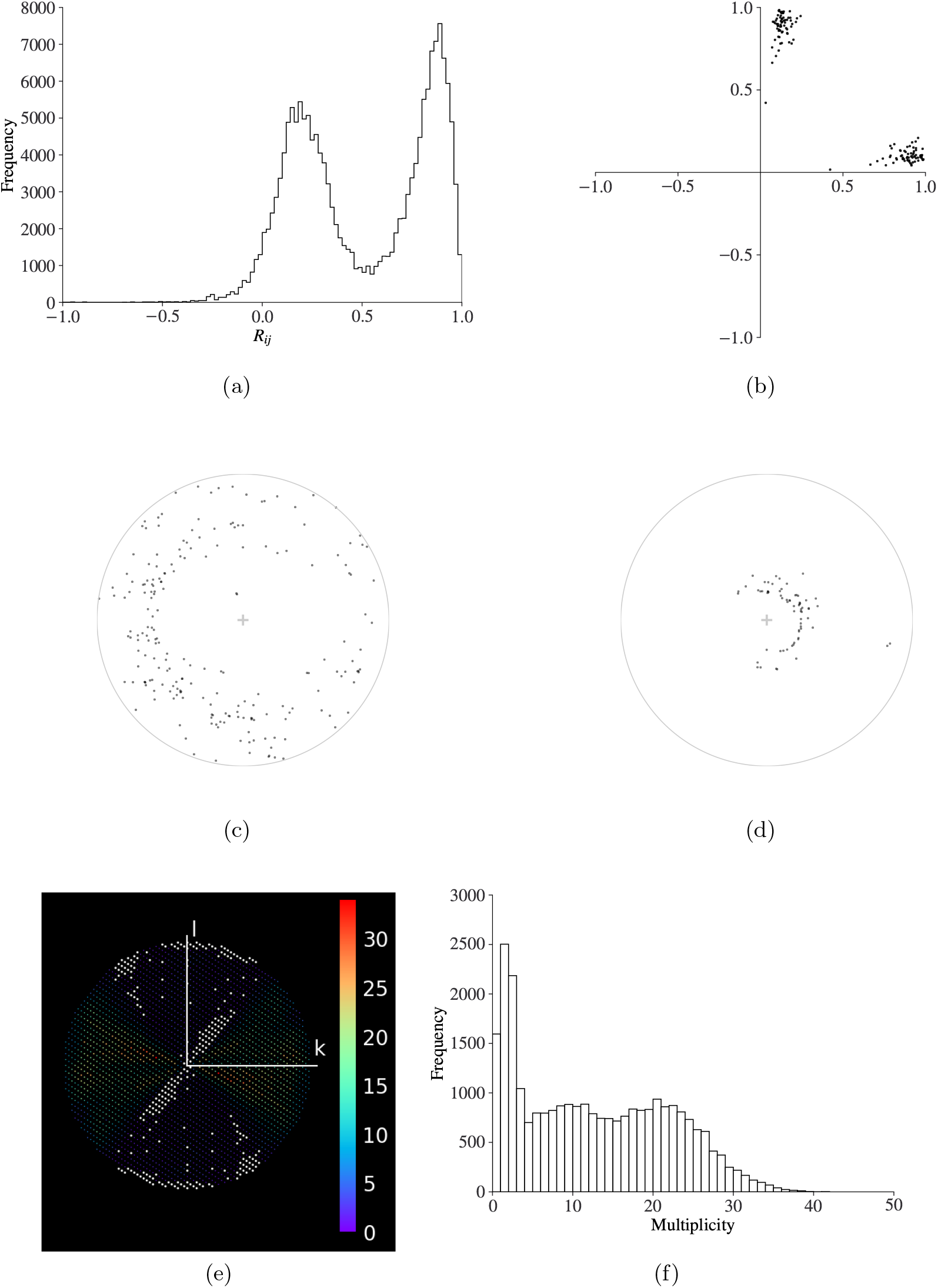
(a) A clear bimodal distribution of the histogram of pairwise *Rij* values is a strong indicator of the presence of an indexing ambiguity. (b) The vectors **x** determined by the minimisation of Equation 2 in dials.cosym. The separation of the vectors into two clusters indicates the presence of an indexing ambiguity. (c) and (d) Stereographic projections of crystal orientations for TehA crystals, representing the direction of *hkl* = 100 and *hkl* = 001 for each crystal respectively, relative to the beam direction (z) which is shown as the central ‘+’ into the page. A point close to the centre of the circle indicates that the crystal axis is close to parallel with the beam, whereas a point close to the edge of the unit circle indicates that the crystal axis is close to perpendicular with the beam. Preferential orientation can lead to regions with systematically low multiplicity or missing reflections. (e) shows the reflection multiplicities in the 0*kl* plane, where white corresponds to missing reflections. The bivariate distribution of multiplicities shown in (f) is also indicative of an uneven distribution of multiplicities.

The best overall unit cell was determined by `dials.two theta refine` as *a* = *b* = 98.76Å, *c* = 136.77Å, and data were scaled together with `dials.scale`. Resolution analysis with `dials.estimate resolution` identified 2.14Å as the resolution where ≈ the fit of a hyperbolic tangent to 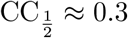.

Six cycles of scaling and filtering were performed by `dials.scale`, where exclusion was performed on whole data sets. A single outlier data set (with a cutoff of 3*σ*) was removed at each of the first five cycles, removing a total of 6.2% of reflections. No significant outliers were identified in the sixth and final cycle.

Structure refinement was performed by REFMAC (Murshudov *et al*., 2011) via *DIMPLE*, using the model from PDB entry 4ycr (Axford *et al*., 2015), using all scale data, and after filtering of outliers using the 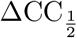 method. Filtering of outlier data sets leads to a slight improvement in merging statistics, particularly in < *I/σ*(*I*) > and *R*_pim_. There is also a slight reduction in the *R*_work_ and *R*_free_ reported by REFMAC.

Stereographic projections of crystal orientation with `dials.stereographic projection` shows that preferential crystal orientatation may be an issue for this experiment (Figures 5c and d). Figures 5e and f show the consequences this has on the distribution of multiplicities in the resulting data set. Analysis with `dials.missing reflections` identifies a single region of missing reflections, comprising 1390 reflections (5.2%) covering the range 53.41 *−* 2.14°A.

## 5. Applications

### 5.1. in situ ligand screening studies of SARS-CoV-2 main protease

With the emergence of the novel coronavirus SARS-CoV-2 and the associated coronavirus disease 2019 (COVID-19), the SARS-CoV-2 main protease has quickly emerged as one of the primary targets for antiviral drug development (Jin *et al*., 2020; Jin *et al*., 2021; Walsh *et al*., 2021). Fragment screening experiments using the XChem platform at Diamond Light Source (Cox *et al*., 2016; Collins *et al*., 2017; Krojer *et al*., 2017) screened over 1250 unique chemical fragments, yielding 74 fragment hits (Douangamath *et al*., 2020).

Fragment screening experiments such as these are typically carried out using conventional cryogenic conditions to minimise the effects of radiation damage, with each structure obtained from a single crystal. Room-temperature data, however, can usefully identify or rule out structural artefacts induced by pushing the temperature far from the biologically relevant level (Durdagi *et al*., 2021; Guven *et al*., 2021).

Over the course of several beamline visits, room-temperature *in situ* data were collected for 30 ligand soaks that were previously shown to bind under cryogenic conditions. Here we highlight room-temperature data collections for five ligand soaks that showed evidence of ligand binding at room-temperature: Z1367324110 (PDB: 5R81) and Z31792168 (PDB: 5R84) (Douangamath *et al*., 2020), Z4439011520 (PDB: 5RH5), Z4439011584 (PDB: 5RH7), and ABT-957 (PDB: 7AEH) (Redhead *et al*., 2021).

Data were collected on beamline I24 at Diamond Light Source, using a DECTRIS PILATUS 3 6M detector, using a 30 *×* 30 µm beam with a flux of approximately 2 *×* 10^11^ photons s^*−*1^. 20^*?*^ of data were collected per crystal with an oscillation range of 0.1^*?*^ and exposure time of 0.02 s per image. Starting angle was varied to maximise total angular range within the constraints imposed by the experimental setup. Based on typical crystal dimensions of 50 µm × 50 µm × 5 µm, X-ray dose per data collection was estimated to be in the range 50 *−*67 kGy using *RADDOSE-3D* (Zeldin *et al*., 2013; Bury *et al*., 2018).

As described in §3, data sets were automatically processed individually with *DIALS* via *xia2*, followed by combined scaling and merging after each data collection with `xia2.multiplex`. Automatic structure refinement and difference map calculations were performed using *DIMPLE*

410 data sets were collected in a single visit at a maximum throughput of 46 data sets per hour. The median time from end of data collection to the completion of the associated processing job was 222.5 s and 352 s for `xia2.multiplex` and *DIMPLE* respectively. 98% of dimple results were reported within 10 minutes of data collection finishing. (See also Supplementary Figure 1.)

Figures 6a and b show the improvement in the merging statistics for the autoprocessed data with the addition of each new data set. There is a visible improvement in the quality of the *DIMPLE* electron density map with the number of crystals (Figures 6e-g).

**Fig. 6.**
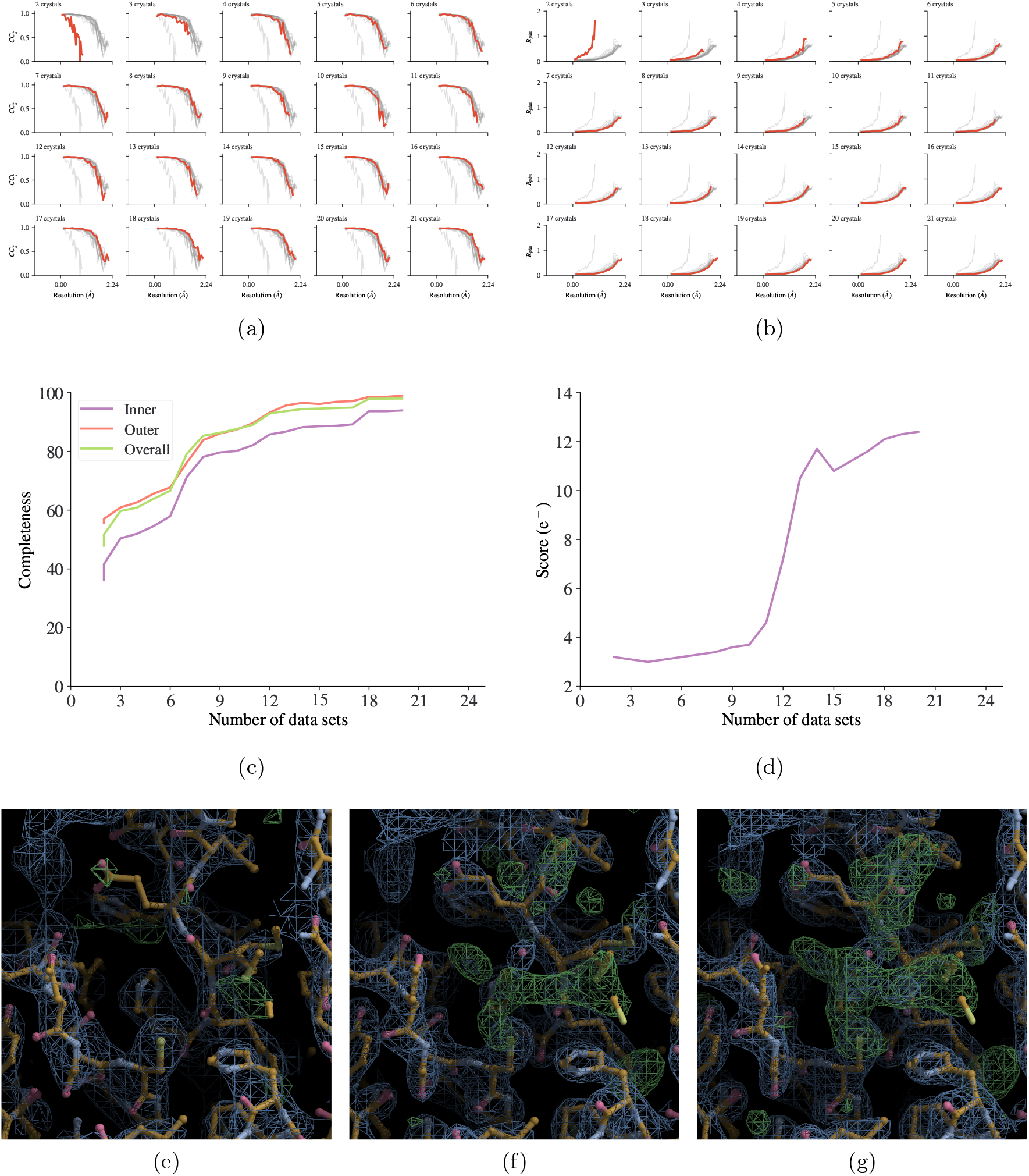
Incremental processing with `xia2.multiplex` and *DIMPLE* on *in situ* data collections of SARS-CoV-2 main protease ligand soak Z4439011520. (a) and (b) CC1*/*2 and *R*_pim_ data processing statistics for ligand Z4439011520 with the inclusion of progressively more data sets, in data collection order, top left to bottom right. and (d) overall data completeness and gemmi (https://gemmi.readthedocs.io) blob search scores. (e), (f) and (g) the ligand density in the autoprocessed *DIMPLE* maps for 2, 9 and 20 crystals respectively. All contours are drawn at 3*σ*.

Analysis of the distribution of unit cell parameters and clustering on unit cell parameters indicated the presence of potential outlier data sets (Figures 7a and b). Reprocessing with a lower unit cell clustering threshold resulted in improved merging statistics for some data sets (Figures 7e and f). Alternatively, 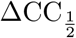 analysis may be useful in identifying outlier data sets. For ligand soak Z4439011520, 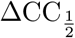 analysis by `dials.scale` identified two outlier data sets over two rounds of scaling and filtering (Figures 7c and d). 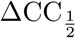 -filtering removed data sets 0 and 18, which were also the two least compatible data sets identified by unit cell clustering, although only the latter was identified as an outlier according to the chosen unit cell clustering threshold. Using the data improved by rejection of outlier data sets as above, initial structure solution was performed using MOLREP (Vagin & Teplyakov, 2010) with 7AEH as the search model. Structures were refined for 200 cycles in REFMAC5 (Murshudov *et al*., 2011) using rigid body refinement, followed by iterative rounds of restrained refinement with automatic TLS and assisted model building in COOT (Emsley *et al*., 2010). Final data processing and refinement statistics for five ligand soaks, Z1367324110, Z31792168, Z4439011520, Z4439011584 and ABT-957, are reported in Table 4. Final coordinates and structure factors have been deposited in the Protein Data Bank (PDB entries 7QT6, 7QT5, 7QT7, 7QT9 and 7QT8 respectively) and raw data uploaded to Zenodo (https://doi.org/10.5281/zenodo.5837942, https://doi.org/10.5281/zenodo.5837946, https://doi.org/10.5281/zenodo.5837903, https://doi.org/10.5281/zenodo.5836055 and https://doi.org/10.5281/zenodo.5837958).

**Table 4.** Data collection, merging and refinement statistics for MPro in situ data sets, after filtering of outliers according to 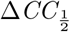. Values in parentheses are for the highest resolution shell.

|  | Z1367324110 | Z31792168 | Z4439011520 | Z4439011584 | ABT-957 |
| --- | --- | --- | --- | --- | --- |
| Data Collection |  |  |  |  |  |
| Exposure time (s) | 0.02 | 0.02 | 0.02 | 0.02 | 0.02 |
| $\Omega$ width ($^{\circ}$) | 0.1 | 0.1 | 0.1 | 0.1 | 0.1 |
| Wavelength ($\text{\AA}$) | 0.9999 | 0.9999 | 0.9999 | 0.9999 | 0.9999 |
| Transmission (%) | 2.9 | 2.9 | 2.9 | 2.9 | 2.9 |
| Number of images | 200 | 200 | 200 | 200 | 200 |
| Number of data sets | 27 | 19 | 19 | 16 | 33 |
| Crystal parameters |  |  |  |  |  |
| Space group | C 1 2 1 | C 1 2 1 | C 1 2 1 | C 1 2 1 | P 1 21 1 |
| Unit-cell parameters ($\text{\AA}$) | $a = 115.21$<br>$b = 54.78$<br>$c = 45.34$<br>$\beta = 101.24$ | $a = 114.77$<br>$b = 54.59$<br>$c = 45.31$<br>$\beta = 101.48$ | $a = 115.69$<br>$b = 54.47$<br>$c = 45.25$<br>$\beta = 101.70$ | $a = 115.86$<br>$b = 54.48$<br>$c = 45.20$<br>$\beta = 101.42$ | $a = 45.23$<br>$b = 54.68$<br>$c = 116.54$<br>$\beta = 100.35$ |
| Data statistics |  |  |  |  |  |
| Resolution range ($\text{\AA}$) | 49.31-2.11<br>(2.19-2.11) | 44.42-2.26<br>(2.34-2.26) | 44.32-2.25<br>(2.33-2.25) | 56.80-2.43<br>(2.52-2.43) | 49.37-2.01<br>(2.08-2.01) |
| No. of unique reflections | 16050 (1586) | 12834 (1277) | 12607 (1272) | 10345 (1038) | 37112 (3748) |
| Multiplicity | 9.8 (9.9) | 7.0 (7.0) | 7.2 (7.3) | 5.9 (5.9) | 12.1 (12.2) |
| $R_{\text{merge}}$ | 0.170 (2.429) | 0.170 (1.956) | 0.162 (1.538) | 0.166 (1.241) | 0.291 (2.409) |
| $R_{\text{meas}}$ | 0.179 (2.551) | 0.184 (2.110) | 0.174 (1.652) | 0.183 (1.380) | 0.304 (2.511) |
| $R_{\text{pim}}$ | 0.053 (0.755) | 0.067 (0.767) | 0.061 (0.577) | 0.073 (0.578) | 0.084 (0.691) |
| Completeness (%) | 99.7 (99.9) | 98.6 (99.6) | 95.2 (97.2) | 97.9 (97.7) | 98.8 (99.8) |
| $\langle I/\sigma(I) \rangle$ | 7.9 (0.6) | 9.9 (1.2) | 8.9 (1.1) | 10.8 (2.0) | 3.6 (0.4) |
| $CC_{\frac{1}{2}}$ | 0.996 (0.331) | 0.994 (0.425) | 0.987 (0.311) | 0.987 (0.305) | 0.992 (0.360) |
| Refinement |  |  |  |  |  |
| $R_{\text{work}}$ | 0.177 | 0.168 | 0.163 | 0.150 | 0.204 |
| $R_{\text{free}}$ | 0.222 | 0.231 | 0.216 | 0.200 | 0.237 |
| R.m.s.d., bond lengths ($\text{\AA}$) | 0.0133 | 0.0107 | 0.107 | 0.0132 | 0.0123 |
| R.m.s.d., bond angles ($^{\circ}$) | 1.843 | 1.691 | 1.81 | 1.903 | 1.752 |
| Average protein $B$ factor ($\text{\AA}^2$) | 55.54 | 52.98 | 50.45 | 48.72 | 36.75 |
| Average water $B$ factor ($\text{\AA}^2$) | 47.13 | 46.09 | 46.44 | 41.28 | 29.5 |
| Average Ligand $B$ factor ($\text{\AA}^2$) | 90.25 | 58.13 | 69.91 | 63.09 | 47.13 |
| Ramachandran (%) |  |  |  |  |  |
| Favoured | 96.69 | 96.04 | 97.35 | 96.36 | 97.03 |
| Allowed | 2.32 | 2.97 | 1.66 | 2.65 | 2.31 |
| PDB Code | 7QT6 | 7QT5 | 7QT7 | 7QT9 | 7QT8 |

**Fig. 7.**
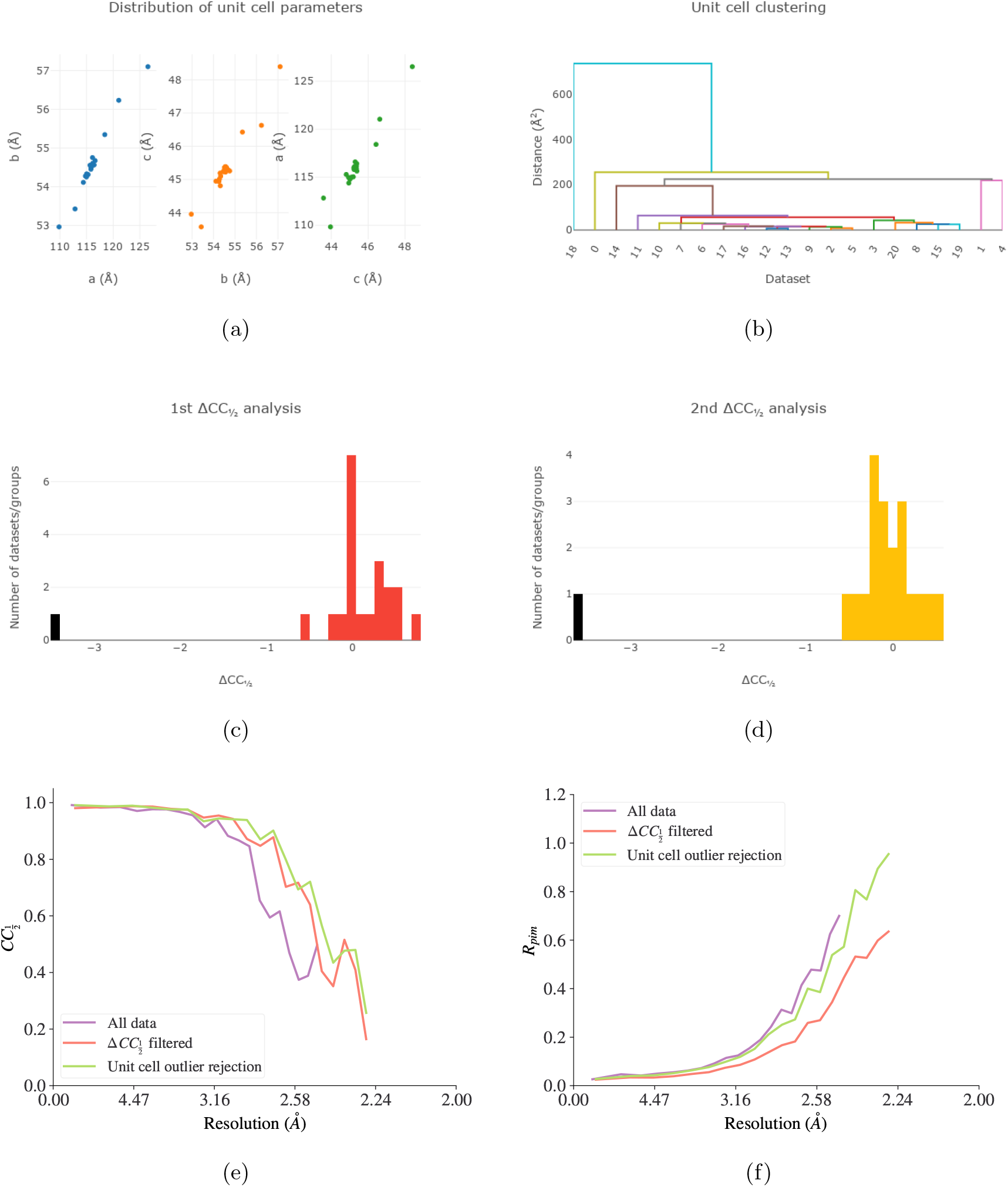
Outlier identification and removal for SARS-CoV-2 main protease ligand soak Z4439011520. Visualisation of the distribution of unit cell parameters (a) and clustering on unit cell parameters (b) may suggest possible outlier data sets. 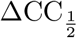- filtering with `dials.scale` can also remove data sets that strongly disagree with the majority of data sets (c) and (d). Removing outlier data sets can improve overall merging statistics (e) and (f).

Ligand soak ABT-957 is of particular interest, as this unexpectedly crystallised in space group *P* 21, in contrast to the space group *C*2 typical of this protein, and indeed observed for the cryo-structure of this ligand (Redhead *et al*., 2021). Autoprocessing (including both *xia2* and `xia2.multiplex`) was performed both using the user-specified target space group, *C*2, and with automatic space group determination. Out of 42 data sets collected, 18 data sets were successfully autoprocessed with *DIALS* via *xia2* in the target space group *C*2, and combined with `xia2.multiplex`. In contrast, all 42 data sets individually processed successfully with automatic space group determination, in a mixture of space groups *P* 1, *P* 2, *P* 21 and *C*2. 33 data sets remained after filtering for inconsistent unit cells. Analysis of symmetry with `dials.cosym` identified the Patterson group *P* 2*/m*, which features an indexing ambiguity due to the approximate pseudo-symmetry of the supergroup *C*2 (Tables 5 and 6).

**Table 5.**
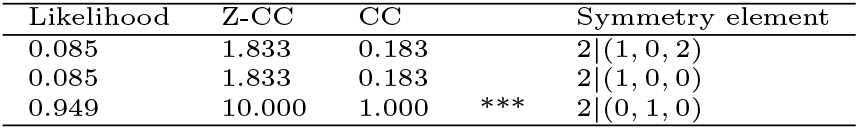
dials.cosym scores for individual symmetry elements for Mpro ligand soak ABT-957

**Table 6.**
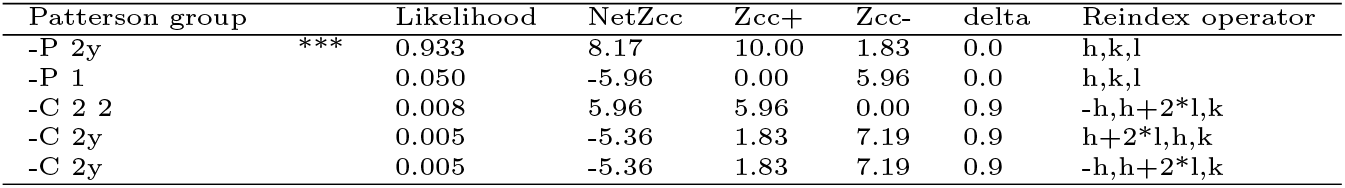
dials.cosym subgroup scores for Mpro ligand soak ABT.

Of the ligand soaked structures collected all showed a near identical binding conformation between cryogenic and room temperature structures. A minor difference was observed in the conformation of ABT-957 with the C9-N-C1(R) amide bond in the room temperature structure being flipped compared to the cryogenic structure (Figure 8). This amide flip had a knock on effect on the rotomer of the gamma-lactam ring and the benzylic side chain which stems from N1 of the gamma lactam.

**Fig. 8.**
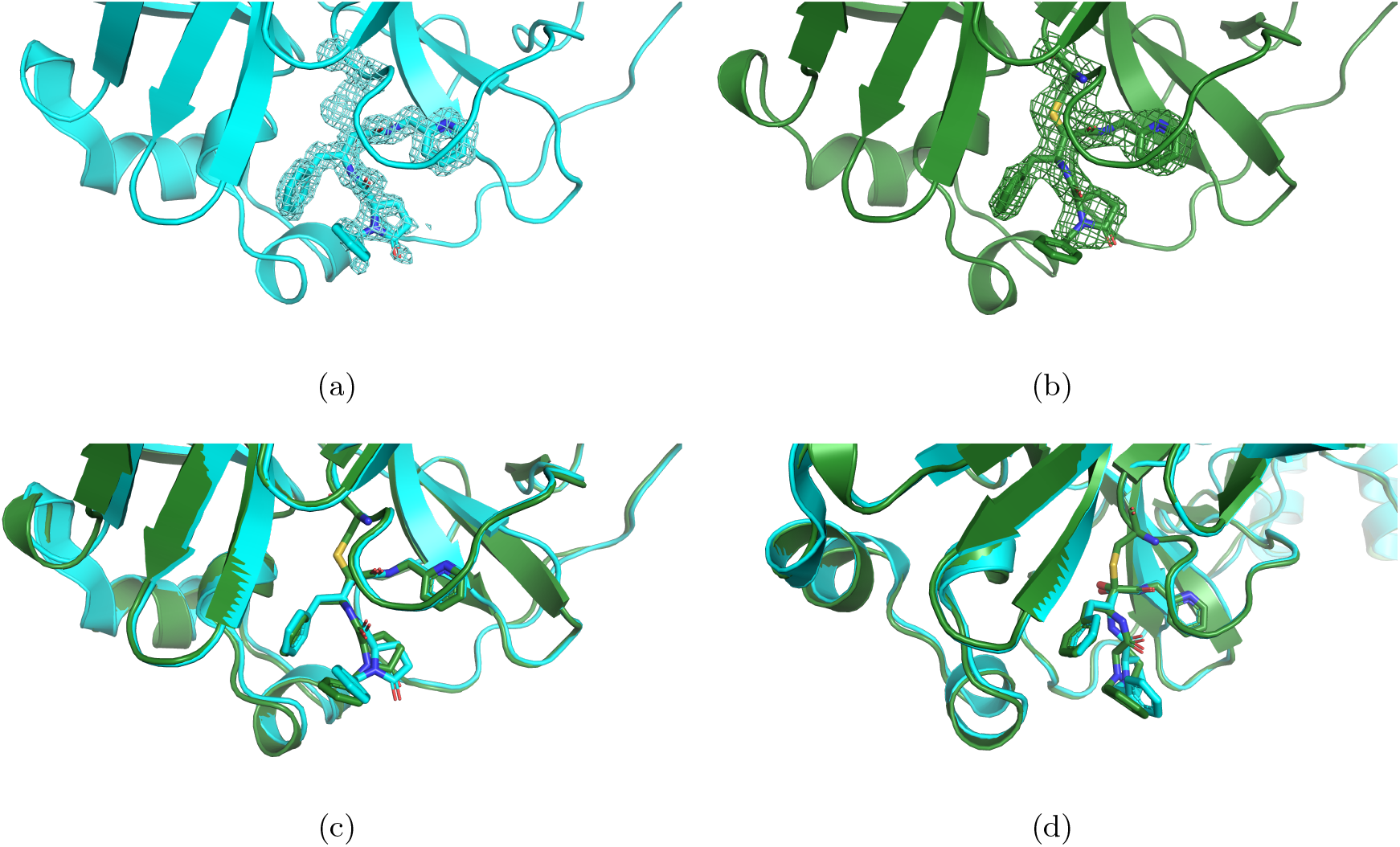
Views of the active site for SARS-CoV-2 main protease (M^pro^) in complex with ABT-957 (a) under cryogenic conditions (Redhead *et al*., 2021) and (b) at room temperature. Contours for the ligand density are drawn at 3*σ*. (c) and (d) two slightly displaced views of the active site for SARS-CoV-2 main protease in complex with ABT-957 to show the conformational differences observed particularly for the oxopyrrolidine and benzyl moieties of ABT-957 when bound to M^pro^at cryo (cyan) and room temperature (green). The structures were superimposed using PyMOL (Schrödinger LLC, 2020).

Inspection of a plot of *R*_*cp*_ vs image number (Supplementary Figure 2) showed slight signs of radiation damage for some ligand soaks. Whilst limiting the number of images used from each data set may lead to improvements to some merging statistics (Supplementary Figure 3), at the cost of completeness and multiplicity, this didn’t lead to any appreciable difference in the ligand density of the final structures (Supplementary Figure 4).

## 6. Conclusions

`xia2.multiplex` has been developed to perform symmetry analysis, scaling and merging of multiple data sets. `xia2.multiplex` is distributed with *DIALS* and hence *CCP4*, and is available as part of the autoprocessing pipelines across MX beamlines at Diamond Light Source, including integration with downstream phasing pipelines such as *DIMPLE* and *Big EP*. It is capable of providing near real-time feedback on data quality and completeness during ongoing multi-crystal data collections, and can be used as part of an iterative workflow to obtain the best possible final data set after an experiment.

We have demonstrated its applicability using two previously-published room-temperature *in situ* multi-crystal data sets, including an example of experimental phasing. Using data sets collected as part of *in situ* room-temperature fragment screening experiments on the SARS-CoV-2 main protease, we have shown the ability of `xia2.multiplex` to provide rapid feedback during multi-crystal experiments, including the identification of an unexpected change in space group with ligand addition.

Remaining challenges include automatic identification of the best subset(s) of data to use for downstream analyses, and providing a user interface via applications such as SynchWeb or *CCP4* to view results and facilitate an interactive workflow using `xia2.multiplex`. Support for MTZ files as input is planned in order to support running `xia2.multiplex` on the output of other data processing software such as *XDS* (Kabsch, 2010) and *MOSFLM* (Battye *et al*., 2011).

## Acknowledgements

The authors would like to thank the authors of DIALS development team for the various components that provide the foundations of `xia2.multiplex`, and those within the wider Diamond Light Source software team who have assisted in the deployment of `xia2.multiplex`. We would also like to thank the Diamond XChem team for their assistance with the SARS-CoV-2 main protease ligand screening experiments, and all MX beamline staff at Diamond Light Source who have provided feedback on `xia2.multiplex` throughout its development.

## Synopsis

A new program, `xia2.multiplex`, has been developed to facilitate symmetry analysis, scaling and merging of multi-crystal data sets.

