## Supporting Information for "xia2.multiplex: a multi-crystal data analysis pipeline"

Richard J. Gildea[a], James Beilsten-Edmands[a], Danny Axford[a], Sam Horrell[a], Pierre Aller[a], James Sandy[a], Juan Sanchez-Weatherby[a], C. David Owen[a,b], Petra Lukacik[a,b], Claire Strain-Damerell[a,b], Robin L. Owen[a], Martin A. Walsh[a,b], and Graeme Winter[a]

[a]Diamond Light Source Ltd, Diamond House, Harwell Science and Innovation Campus, Didcot, Oxfordshire, OX11 0DE, UK
[b]Research Complex at Harwell, Harwell Science and Innovation Campus, Didcot, OX11 0FA, UK

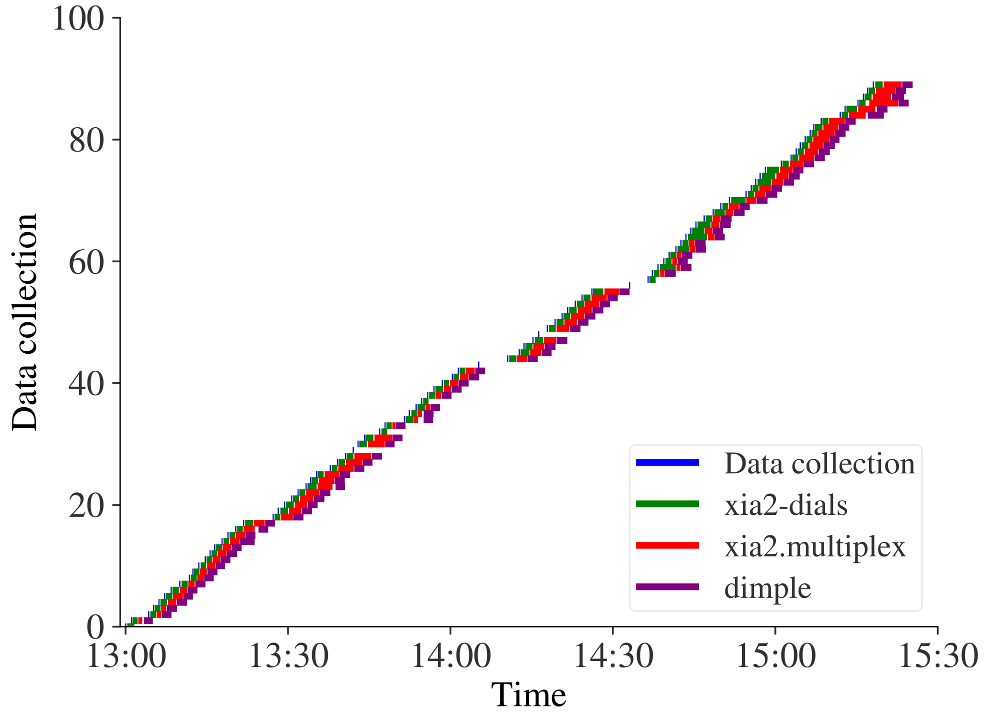


Figure 1: Real-time multi-crystal autoprocessing of a subset of the SARS-CoV-2 main protease data collections reported in §5, using *xia2*, *DIALS*, `xia2.multiplex` and *DIMPLE*. 410 data sets were collected in a single visit at a maximum throughput of 46 data sets per hour. The median time from end of data collection to the completion of the associated processing job was 222.5 s and 352 s for `xia2.multiplex` and *DIMPLE* respectively. 98% of dimple results were reported within 10 minutes of data collection finishing.

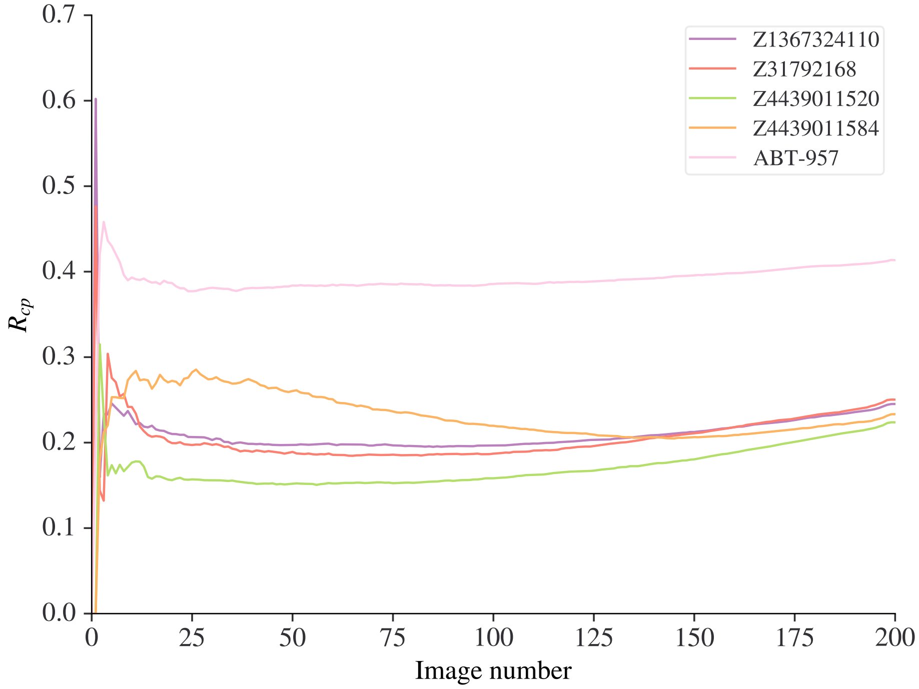


Figure 2: $R_{cp}$ vs image number for the SARS-CoV-2 main protease data collections reported in §5. This suggests some signs of slight radiation damage after around 100 images for the Z1367324110, Z31792168 and Z4439011520 data sets.

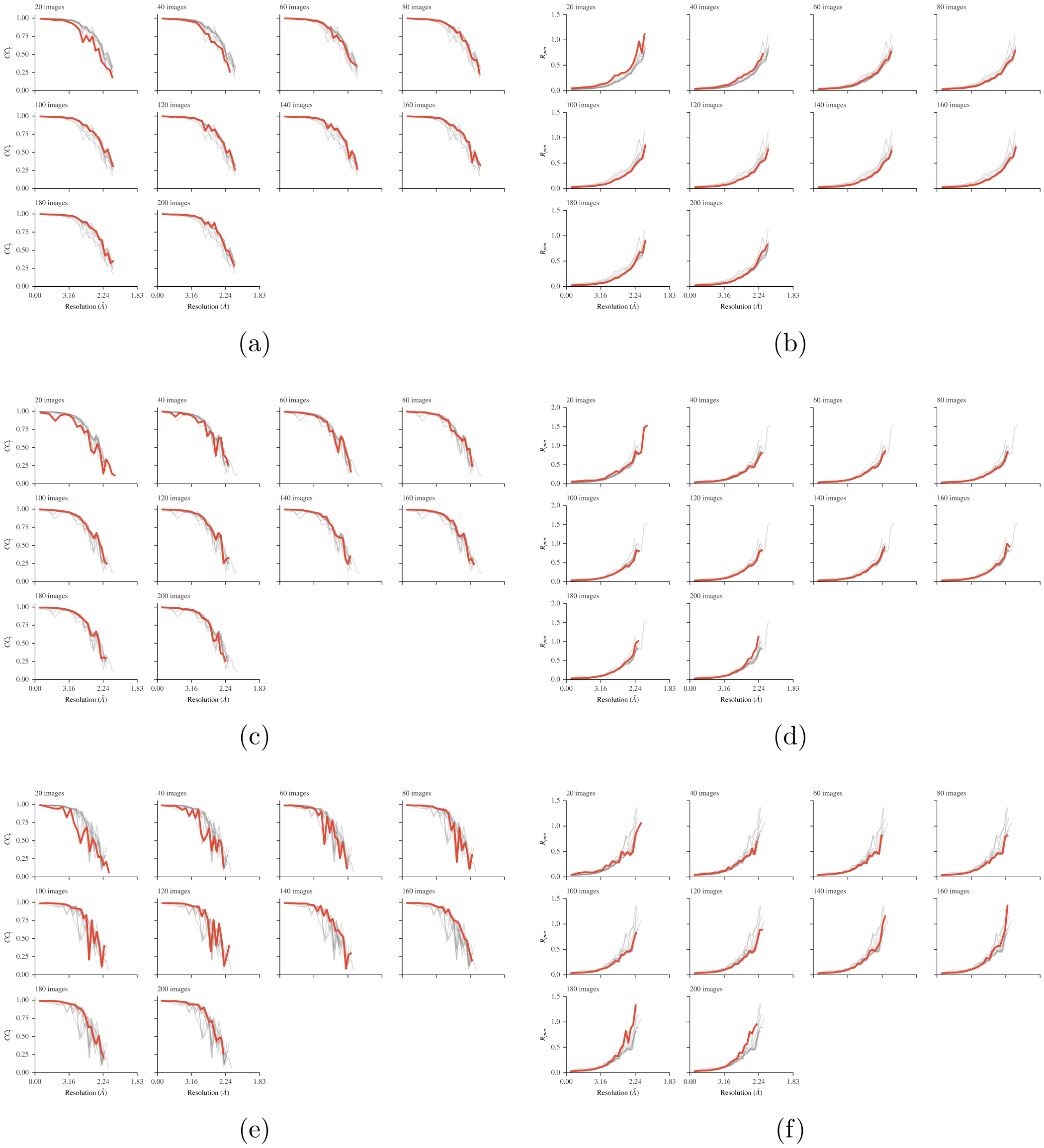


Figure 3: Comparison of merging statistics using only the first 20, 40, ..., 200 images from each data set for SARS-CoV-2 main protease ligand soaks Z1367324110 (a) and (b), Z31792168 (c) and (d) and Z4439011520 (e) and (f).

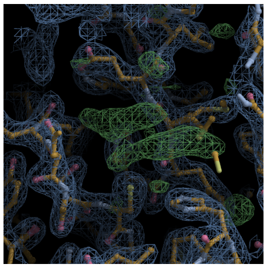
(a)

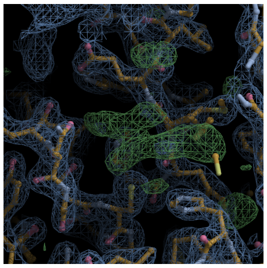
(b)

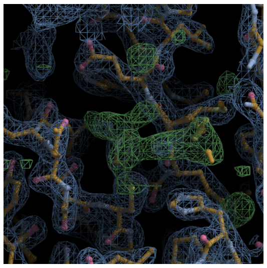
(c)

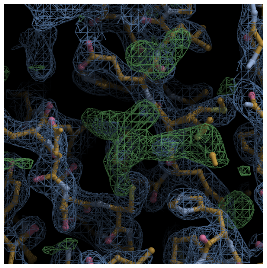
(d)

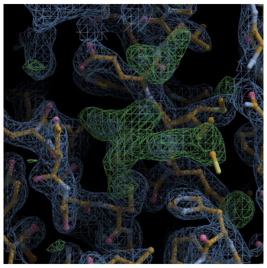
(e)

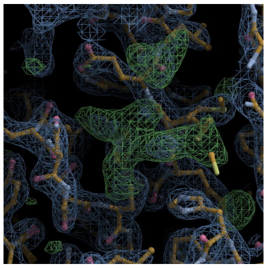
(f)

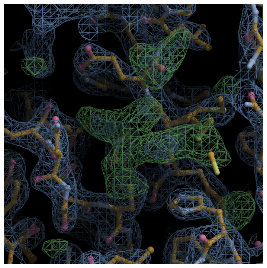
(g)

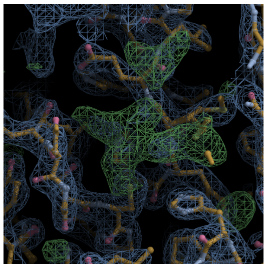
(h)

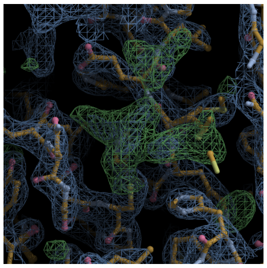
(i)

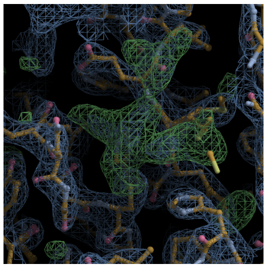
(j)

Figure 4: *DIMPLE* difference electron density for SARS-CoV-2 main protease ligand soak Z4439011520 using only the first 20 (a), 40 (b), ..., 200 (j) images of each data set. All contours are drawn at $3\sigma$.

### Dose calculations

RADDOSE-3D input and output for dose calculation for a typical crystal as used in §5 hit side-on with the X-ray beam:

#### Input

```
##############################################################################
# Crystal Block #
##############################################################################

Crystal

Type Cuboid
# Crystal shape can be Cuboid or Spherical

Dimensions 5 50 50
# Dimensions of the crystal in X,Y,Z in µm.
# Z is the beam axis, Y the rotation axis and
# X completes the right handed set
# (vertical if starting face-on).

PixelsPerMicron 0.5
# This defines the coarseness of the simulation
# (i.e. how many voxels the crystal is divided into.)
# Preferably set as high as possible, however for a higher
# value the simulation will take longer to complete.
# Recommended to try increasing between 0.5 and 5 and ensure
# the reported dose value converges as PixelsPerMicron increases.
# As a rule of thumb, this needs to be at least 10x the beam
# FWHM for a Gaussian beam.
# e.g. 20 µm FWHM beam -> 2 µm voxels -> 0.5 voxels/µm

# NOTE: Use AngleP/AngleL if your crystal is not face-on to the beam.
# See RD3D user guide for more details

# Also need to specify the crystal composition below (Example case for insulin given):
AbsCoefCalc RD3D
# Absorption Coefficients calculated
# using RADDOSE-3D (Zeldin et al. 2013).

UnitCell 115.21 54.78 45.34 90 101.24 90
# unit cell size: a, b, c with alpha, beta and gamma angles default to 90°

NumMonomers 2
# number of monomers in unit cell

NumResidues 305
# number of residues per monomer

ProteinHeavyAtoms S 22
# heavy atoms added to protein part of the
# monomer, i.e. S, coordinated metals, Se in Se-Met

SolventHeavyConc S 700
```

```
# concentration of elements in the solvent
# in mmol/l. Oxygen and lighter elements
# should not be specified

SolventFraction 0.3716
# fraction of the unit cell occupied by solvent

##############################################################################
# Beam Block #
##############################################################################

Beam

Type Gaussian
# beam profile can be Gaussian or TopHat
# Flux 7e12
Flux 2.03e11
# in photons per second (2e12 = 2 * 10^12)
FWHM 30 30
# in µm, horizontal by vertical for a Gaussian beam
Energy 12.4
# photon energy in keV

Collimation Rectangular 100 100
# Horizontal/Vertical collimation of the beam
# For 'uncollimated' Gaussians, 3xFWHM recommended

##############################################################################
# Wedge Block #
##############################################################################

Wedge 0 20
# Start and End rotational angle of the crystal with Start < End

ExposureTime 2
# Total time for entire angular range

# AngularResolution 2
# Only change from the defaults when using very
# small wedges, e.g 5°.

# NOTE: To define more complex geometries (helical, de-centred, or offset),
# see the StartOffset, TranslatePerDegree, and RotAxBeamOffset keywords
# in the User Guide
```

#### Output

```
Cuboid (Polyhedron) crystal of size [5, 50, 50] um [x, y, z] at a resolution of 2.00 microns per voxel edge.
Simple DDM.
Gaussian beam, 100.0x100.0 um with 30.00 by 30.00 FWHM (x by y) and 2.0e+11 photons per second at 12.40 keV.
Wedge 1:
Collecting data for a total of 2.0s from phi = 0.0 to 20.0 deg.

Crystal coefficients calculated with RADDOSE-3D.
Photelectric Coefficient: 1.83e-04 /um.
Inelastic Coefficient: 1.29e-05 /um.
Elastic Coefficient: 1.36e-05 /um.
Attenuation Coefficient: 2.09e-04 /um.
Density: 0.78 g/ml.

Average Diffraction Weighted Dose         : 0.066779 MGy
Last Diffraction Weighted Dose            : 0.126371 MGy
Elastic Yield                             : 4.73e+07 photons
Diffraction Efficiency (Elastic Yield/DWD): 7.09e+08 photons/MGy
Average Dose (Whole Crystal)              : 0.108026 MGy
Average Dose (Exposed Region)             : 0.108026 MGy
Max Dose                                  : 0.183466 MGy
Average Dose (95.0 % of total absorbed energy threshold (0.05 MGy)): 0.121631 MGy
Dose Contrast (Max/Threshold Av.)         : 1.51
Used Volume                               : 100.0%
Absorbed Energy (this Wedge)              : 1.30e-06 J.
Dose Inefficiency (Max Dose/mJ Absorbed)  : 141.3 1/g
Dose Inefficiency PE (Max Dose/mJ Deposited): 144.4 1/g
Final Dose Histogram:
Bin  1,  0.0 to  0.1 MGy: 44.7 %
Bin  2,  0.1 to  3.4 MGy: 55.3 %
Bin  3,  3.4 to  6.7 MGy:  0.0 %
Bin  4,  6.7 to 10.1 MGy:  0.0 %
Bin  5, 10.1 to 13.4 MGy:  0.0 %
Bin  6, 13.4 to 16.7 MGy:  0.0 %
Bin  7, 16.7 to 20.0 MGy:  0.0 %
Bin  8, 20.0 to 23.4 MGy:  0.0 %
Bin  9, 23.4 to 26.7 MGy:  0.0 %
Bin 10, 26.7 to 30.0 MGy:  0.0 %
Bin 11, 30.0 MGy upwards:  0.0 %
```

RADDOSE-3D input and output for dose calculation for a typical crystal as used in §5 hit face-on with the X-ray beam:

#### Input

```
##############################################################################
# Crystal Block #
##############################################################################

Crystal

Type Cuboid
# Crystal shape can be Cuboid or Spherical

Dimensions 50 50 5
# Dimensions of the crystal in X,Y,Z in µm.
# Z is the beam axis, Y the rotation axis and
# X completes the right handed set
# (vertical if starting face-on).

PixelsPerMicron 0.5
# This defines the coarseness of the simulation
# (i.e. how many voxels the crystal is divided into.)
# Preferably set as high as possible, however for a higher
# value the simulation will take longer to complete.
# Recommended to try increasing between 0.5 and 5 and ensure
# the reported dose value converges as PixelsPerMicron increases.
# As a rule of thumb, this needs to be at least 10x the beam
# FWHM for a Gaussian beam.
# e.g. 20µm FWHM beam -> 2µm voxels -> 0.5 voxels/µm

# NOTE: Use AngleP/AngleL if your crystal is not face-on to the beam.
# See RD3D user guide for more details

# Also need to specify the crystal composition below (Example case for insulin given):
AbsCoefCalc RD3D
# Absorption Coefficients calculated
# using RADDOSE-3D (Zeldin et al. 2013).

UnitCell 115.21 54.78 45.34 90 101.24 90
# unit cell size: a, b, c with alpha, beta and gamma angles default to 90°

NumMonomers 2
# number of monomers in unit cell

NumResidues 305
# number of residues per monomer

ProteinHeavyAtoms S 22
# heavy atoms added to protein part of the
# monomer, i.e. S, coordinated metals, Se in Se-Met

SolventHeavyConc S 700
# concentration of elements in the solvent
# in mmol/l. Oxygen and lighter elements
```

```
# should not be specified

SolventFraction 0.3716
# fraction of the unit cell occupied by solvent

##############################################################################
# Beam Block #
##############################################################################

Beam

Type Gaussian
# beam profile can be Gaussian or TopHat
# Flux 7e12
Flux 2.03e11
# in photons per second (2e12 = 2 * 10^12)
FWHM 30 30
# in µm, horizontal by vertical for a Gaussian beam
Energy 12.4
# photon energy in keV

Collimation Rectangular 100 100
# Horizontal/Vertical collimation of the beam
# For 'uncollimated' Gaussians, 3xFWHM recommended

##############################################################################
# Wedge Block #
##############################################################################

Wedge 0 20
# Start and End rotational angle of the crystal with Start < End

ExposureTime 2
# Total time for entire angular range

# AngularResolution 2
# Only change from the defaults when using very
# small wedges, e.g 5°.

# NOTE: To define more complex geometries (helical, de-centred, or offset),
# see the StartOffset, TranslatePerDegree, and RotAxBeamOffset keywords
# in the User Guide
```

#### Output

```
Cuboid (Polyhedron) crystal of size [50, 50, 5] um [x, y, z] at a resolution of 2.00 microns per voxel edge.
Simple DDM.
Gaussian beam, 100.0x100.0 um with 30.00 by 30.00 FWHM (x by y) and 2.0e+11 photons per second at 12.40 keV.
Wedge 1:
Collecting data for a total of 2.0s from phi = 0.0 to 20.0 deg.

Crystal coefficients calculated with RADDOSE-3D.
Photelectric Coefficient: 1.83e-04 /um.
Inelastic Coefficient: 1.29e-05 /um.
Elastic Coefficient: 1.36e-05 /um.
Attenuation Coefficient: 2.09e-04 /um.
Density: 0.78 g/ml.

Average Diffraction Weighted Dose         : 0.050366 MGy
Last Diffraction Weighted Dose            : 0.095817 MGy
Elastic Yield                             : 2.02e+07 photons
Diffraction Efficiency (Elastic Yield/DWD): 4.01e+08 photons/MGy
Average Dose (Whole Crystal)              : 0.069056 MGy
Average Dose (Exposed Region)             : 0.069056 MGy
Max Dose                                  : 0.183705 MGy
Average Dose (95.0 % of total absorbed energy threshold (0.03 MGy)): 0.082169 MGy
Dose Contrast (Max/Threshold Av.)         : 2.24
Used Volume                               : 100.0%
Absorbed Energy (this Wedge)              : 5.53e-07 J.
Dose Inefficiency (Max Dose/mJ Absorbed)  : 332.0 1/g
Dose Inefficiency PE (Max Dose/mJ Deposited): 339.2 1/g
Final Dose Histogram:
Bin  1,  0.0 to  0.1 MGy: 74.2 %
Bin  2,  0.1 to  3.4 MGy: 25.8 %
Bin  3,  3.4 to  6.7 MGy:  0.0 %
Bin  4,  6.7 to 10.1 MGy:  0.0 %
Bin  5, 10.1 to 13.4 MGy:  0.0 %
Bin  6, 13.4 to 16.7 MGy:  0.0 %
Bin  7, 16.7 to 20.0 MGy:  0.0 %
Bin  8, 20.0 to 23.4 MGy:  0.0 %
Bin  9, 23.4 to 26.7 MGy:  0.0 %
Bin 10, 26.7 to 30.0 MGy:  0.0 %
Bin 11, 30.0 MGy upwards:  0.0 %
```
